# Integrating Protein Localization with Automated Signaling Pathway Reconstruction

**DOI:** 10.1101/609149

**Authors:** Ibrahim Youssef, Jeffrey Law, Anna Ritz

## Abstract

Understanding cellular responses via signal transduction is a core focus in systems biology. Tools to automatically reconstruct signaling pathways from protein-protein interactions (PPIs) can help biologists generate testable hypotheses about signaling. However, automatic reconstruction of signaling pathways suffers from many interactions with the same confidence score leading to many equally good candidates. Further, some reconstructions are biologically misleading due to ignoring protein localization information. We propose *LocPL*, a method to improve the automatic reconstruction of signaling pathways from PPIs by incorporating information about protein localization in the reconstructions. The method relies on a dynamic program to ensure that the proteins in a reconstruction are localized in cellular compartments that are consistent with signal transduction from the membrane to the nucleus. *LocPL* and existing reconstruction algorithms are applied to two PPI networks and assessed using both global and local definitions of accuracy. *LocPL* produces more accurate and biologically meaningful reconstructions on a versatile set of signaling pathways. *LocPL* is a powerful tool to automatically reconstruct signaling pathways from PPIs that leverages cellular localization information about proteins. The underlying dynamic program and signaling model are flexible enough to study cellular signaling under different settings of signaling flow across the cellular compartments.

## 1 Introduction

A fundamental goal of molecular systems biology is to understand how individual proteins and their interactions may contribute to a larger cellular response. Repositories for experimentally derived or manually curated human protein-protein interaction (PPI) information [1–7] have been critical for achieving that goal. These databases conceptualize the interaction information as a graph, or an *interactome*, where edges connect proteins that are known to interact. Such interactomes are useful for studying the topology of signaling pathways by forming static networks and focusing on the interconnections between proteins and how signals flow between them. In particular, interaction data have enabled the development of methods that aim to link extracellular signals to downstream cellular responses.

Most methods that link signals with responses were initially applied to yeast studies [8–10]. A handful of the initial methods were applied to human signaling, including the apoptosis pathway [11] and the immune response network [12]. Approaches for identifying relevant static sub-networks have drawn on different graph theoretic methods, including shortest paths [13, 14], Steiner trees and related formulations [15, 16], network flow [9, 17] and random walk approaches [18–20].

As the wealth of PPI information has grown, these methods have been increasingly adopted to study human signaling. PathLinker is a recent pathway reconstruction approach that returns ranked paths for a specific human signaling pathway of interest [13]. Given a weighted interactome, a set of known receptors, and a set of known transcriptional regulators (TRs), PathLinker returns the *k*-shortest paths from any receptor to any transcriptional regulator, and the collection of these paths constitute a *pathway reconstruction*. PathLinker reconstructions have been shown to outperform other pathway reconstruction methods on human networks [13]. PathLinker predicted that CFTR, a chloride ion channel transporter, was involved in Wnt signaling; RNAi and Co-immunoprecipitation experiments confirmed CFTR’s involvement in Wnt signaling in HEK293 cells [13].

### Pathway Reconstruction Challenges

Despite PathLinker’s success, the problem of identifying accurate pathway reconstructions remains challenging. PathLinker paths are prioritized by their reconstruction scores that are the product of a path edge weights. These paths combined form a *pathway reconstruction*. We assessed PathLinker reconstructions for four well-studied and diverse signaling pathways: the Wnt pathway is critical for the development of tissues cell fate specification [21]; the Interleukin-2 (IL2) pathway plays a major role in controlling the immune system and regulating homeostasis [22]; the *α*6*β*4 Integrin pathway regulates cell adhesion to the extracellular matrix [23], and the Epidermal Growth Factor Receptor (EGFR1) pathway regulates cell proliferation, survival, and migration [24]. Careful analysis of the ranked paths across these pathways revealed two main challenges in pathway reconstruction.

First, we found that many PathLinker paths have identical reconstruction scores. For example, about 52% of the paths in the Wnt reconstruction had the same score. This feature was not unique to Wnt; 64%, 82.6%, and 48.2% of the paths were tied in the IL2, *α*6*β*4 Integrin, and EGFR1 pathways, respectively. Strikingly, even the top-ranked paths in the reconstructions were often tied (top 38 paths in Wnt, top 87 paths in IL2, top 57 paths in *α*6*β*4 Integrin, and top 330 paths in EGFR1). We found that the tied paths were a result of many interactions with identical weights in the underlying interactome (Figure 1). For example, in the PathLinker interactome (*PLNet*_1_), nearly 68% of the interactions have only two distinct weight values. In the interactome used in this work (*PLNet*_2_), around 71% of the interactions have just three different weight values. The coarse interaction weighting is also apparent in the HIPPIE network [2], where 55% of the interactions share the same edge weight (Figure 1).

**Figure 1:**
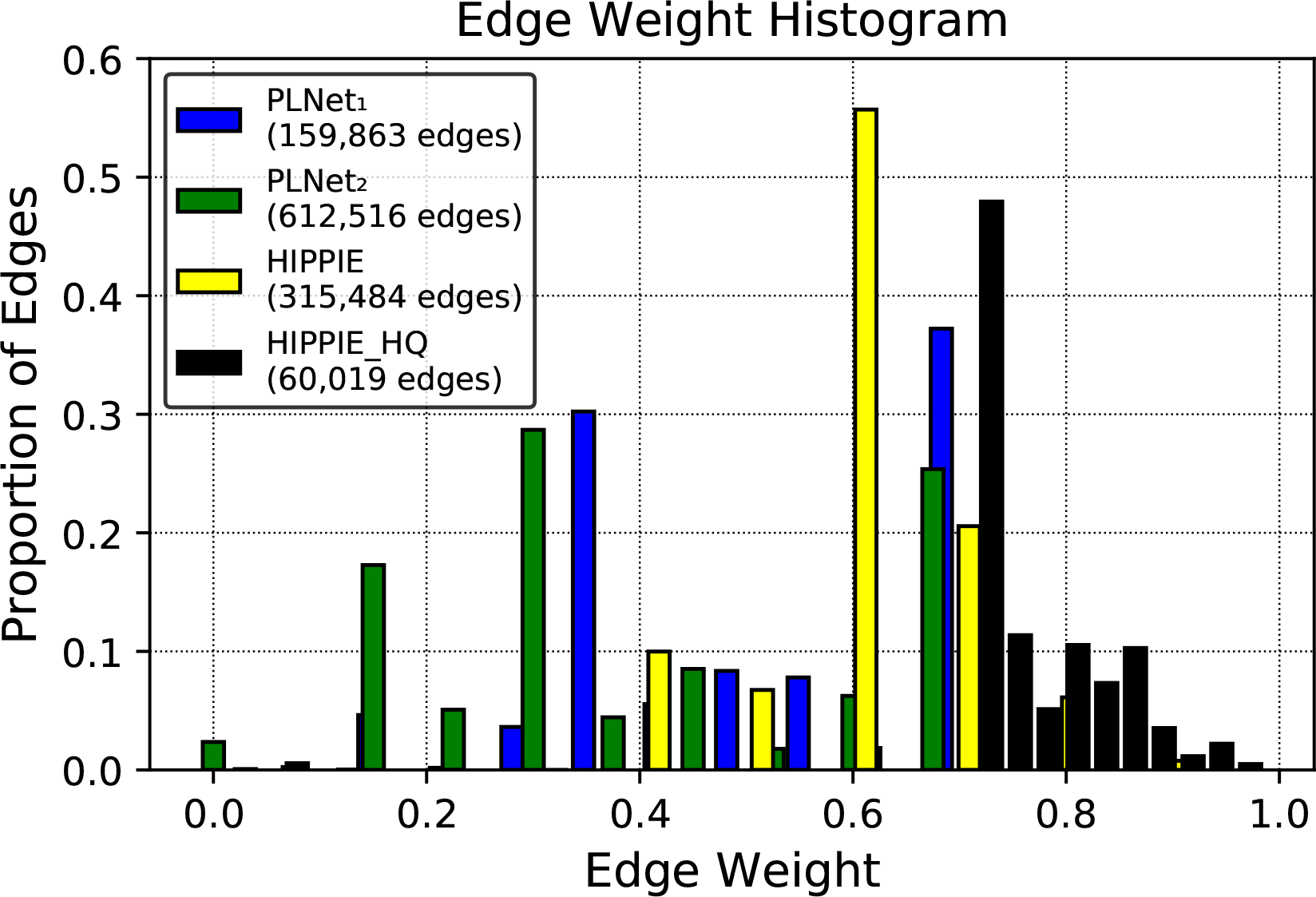
Proportion of edges with identical edge weights in the PathLinker and HIPPIE interactomes. *PLNet*_1_ is the PathLinker interactome [13], while *PLNet*_2_ is the interactome used in this work. The HIPPIE High Quality (HIPPIE HQ) interactome includes all HIPPIE edges with a weight *≥* 0.73 [2]. The histogram number of bins is 10 with a size of 0.02 for each.

Second, we noted that paths in the reconstructions contained a mix of pathway-specific signaling interactions relevant to the pathway under study (*positive interactions*) and non-pathway interactions (we will call them *negative interactions*, though they may very well be signaling interactions relevant to other pathways or pathway-specific interactions that have not been annotated yet). Paths are rarely comprised solely of positive interactions: in all four pathway reconstructions, over 95% of the paths that include at least one positive interaction also contain a negative interaction. PathLinker does not consider protein localization in the pathway reconstructions, so interactions within the same path may be unrealistic in terms of compartment co-localization. Given the first challenge of coarse interaction weights, additional evidence about protein localization could be useful for breaking tied path scores.

To overcome the challenges described above, we sought to incorporate an independent data type into the pathway reconstruction problem. While many methods have integrated gene expression data in pathway reconstructions [15, 9, 20], we wish to improve “canonical” pathways that are independent of a specific context (e.g. a condition or disease). Instead, we make use of information about a protein’s localization within the cell to constrain the paths in a reconstruction.

### Contributions

We propose *LocPL*, an extended version of PathLinker that reconstructs pathways by incorporating information about cellular localization in two ways. First, *LocPL* uses localization information to discard likely false positive interactions from the interactome before running PathLinker, improving its specificity. Second, *LocPL* incorporates the localization information in a dynamic programming scheme to identify spatially-coherent paths and re-prioritize tied paths (Figure 2A). We show that paths with larger proportions of signaling interactions will be promoted higher in the *k*-shortest paths list, and those of smaller proportions will be demoted. We compare the *LocPL* pathway reconstructions to those from PathLinker on two interactomes: a new interactome, *PLNet*_2_, which quadruples the number of interactions compared to the PathLinker interactome, and the HIPPIE interactome [2]. We also compare *LocPL* to a color-coding method [25, 26]. In addition to performing a global performance assessment of paths, we present a local measure to assess path quality individually. Visual inspection of the top 100 paths in the Wnt, IL2, *α*6*β*4 Integrin, and EGFR1 pathway reconstructions reveal that the spatially-coherent approach changes the reconstruction topology, in some cases removing paths that lead to activation of other pathways. This work demonstrates that incorporating protein localization information into signaling pathway reconstruction improves predictions that are necessary for appropriate hypothesis generation.

**Figure 2:**
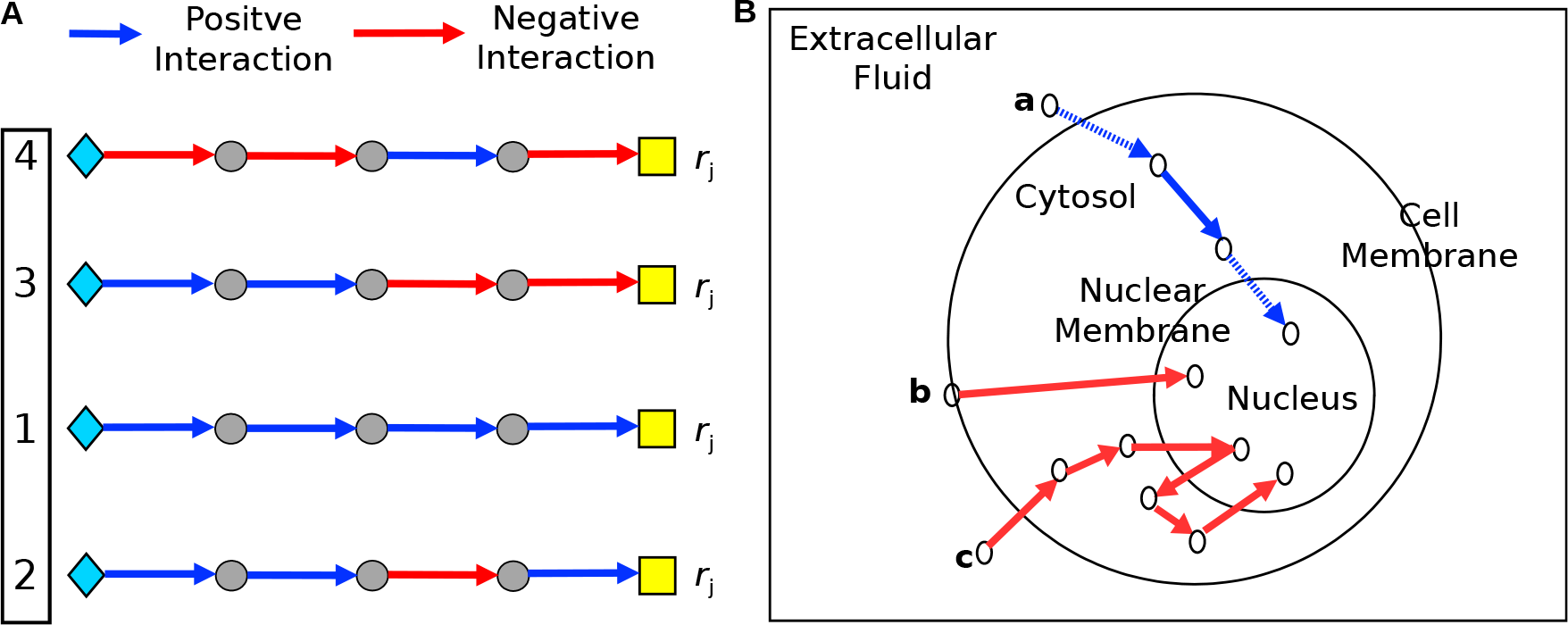
**A.** Illustration of four PathLinker paths from receptors (diamonds) to transcriptional regulators (yellow boxes) that all have the same reconstruction score *r*_*j*_. Blue edges represent true positive interactions, and red edges represent false positives. The goal of breaking ties is to re-rank the tied paths so paths with more positives are ranked higher (black box). **B.** Simplified model diagram for the signaling flow structure. Blue edges represent valid interactions. The blue solid edges are between pairs of proteins sharing one cellular compartment, and the blue dotted edges are proteins that traverse between two compartments. Paths that violate our signaling model assumptions are shown in red, where path (b) has a single interaction between a pair of proteins without a common cellular compartment, and signaling in path (c) does not reside in the nucleus once it reached the nuclear compartment.

## 2 Methods

We first introduce ComPPI, the protein localization database that *LocPL* uses to refine pathway reconstructions, and then we present an overview of *LocPL*. After describing the model used for signaling flow, we present a dynamic program for computing scores that reflect a path’s consistency with the model of signaling. Then, we describe the color-coding method that *LocPL* is compared to. Finally, we detail the interactome and signaling pathway datasets and the means of assessing pathway reconstruction performance.

### 2.1 Localized Protein-Protein Interactions from ComPPI

ComPPI is a database that predicts cellular compartments for human proteins and PPIs [27] (Version 2.1.1, September 10^*th*^, 2018 [28]). For each protein, ComPPI computes *localization scores* describing the likelihood of a protein to be found in one of the major six subcellular compartments: (i) extracellular fluid, (ii) cell membrane, (iii) cytosol, (iv) nucleus, (v) secretory pathway (e.g. transport vesicles), and (vi) mitochondria. ComPPI uses three types of information to infer the localization scores: experimental verification, computational prediction, and unknown sources, resulting in high, medium, and low localization scores, respectively. The *interaction score*, computed by ComPPI from localization scores of the participating proteins, represents the probability that an interaction takes place inside the cell.

### 2.2 *LocPL*: Localized PathLinker

Signaling pathway analysis methods typically take an interactome as input, represented as a graph *G* = (*V, E*) where the nodes *V* are proteins and the edges *E* are PPIs. In the case of *LocPL*, the graph is directed, each edge (*u, v*) *∈ E* has a weight *w*_*uv*_ ∈ [0,1], and every interaction is predicted to occur within some cellular compartment according to ComPPI. *LocPL* uses the ComPPI database to restrict the interactions of the interactome by removing edges with an interaction score of zero – these interactions could take place from a biophysical perspective, but are less likely to occur within the cell due to the predicted protein localization. After this filtration step, all edges in the interactome have a non-zero probabilistic score aggregated across all cellular compartments. For subsequent steps of *LocPL*, we use the ComPPI localization scores that reflect individual proteins in specific cellular compartments.

*LocPL*’s core method is a *k*-shortest path algorithm previously described as PathLinker [13]. Given a directed, weighted interactome *G*, a set *R* of receptors and a set *T* of transcriptional regulators (TRs) for a pathway of interest, and a number of paths *k*, PathLinker outputs a ranked list of the *k* shortest paths, 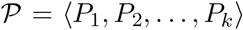, where a path *P*_*i*_ = (*v*_1_*, v*_2_, …, *v*_*m*_) is comprised of *m* nodes that begin at a receptor (*v*_1_ *∈ R*) and ends at a TR (*v*_*m*_ ∈ *T*). Each path *P*_*i*_ is ranked by the product of its edge weights (its *reconstruction score r*_*i*_), and *r*_*i*_ ≥ *r*_*i* +1_ for every *i*. Note that the shortest path is the one whose edge weights product is the highest among all paths since PathLinker takes the negative log-transform of the edge weights at the reconstruction step.

After running PathLinker on the interactome, *LocPL* breaks ties in the candidate list of paths 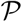 by considering a model of signaling flow based on cellular compartments. For each path *P*_*i*_, a dynamic program identifies the *signaling score s*_*i*_ of the most likely series of compartments for each node that is consistent with the signaling flow model. After this step, each path *P*_*i*_ will have two scores: a reconstruction score *r*_*i*_ computed by PathLinker and a signaling score *s*_*i*_ computed by the dynamic program. The signaling score is used to re-prioritize the tied reconstruction scores by partitioning the paths into ties (e.g. all paths with the same reconstruction score) and reordering the paths within each group in decreasing order of the signaling score (Figure 2A).

### 2.3 Signaling Flow Structure and Assumptions

In order to use protein localization information in pathway reconstructions, we first state some assumptions about the pathways we aim to reconstruct. First, we only consider intracellular signaling that begins with activation of a membrane-bound protein receptor and is transmitted to a DNA-binding transcription factor through PPIs within the cytosol. Hence, we focus on three cellular compartments: a combination of extracellular fluid and cell membrane (*ExtMem*), which represents where a receptor may be located, *Cytosol*, and *Nucleus*. Second, we assume a unidirectional signaling flow from *ExtMem* through *Cytosol* to *Nucleus*. Third, multiple interactions may occur within the same cellular compartment (e.g. multiple interactions may occur within *Cytosol*). Fourth, signaling flow advances through either interacting proteins that share the same cellular compartment, or a protein that can traverse different cellular compartments. These assumptions impose an ordering on the compartments that must be visited, which we will use in breaking tied paths. Figure 2B illustrates these assumptions with three different paths as examples of valid and invalid paths/interactions. Path **a** is valid; however, path **b** is not valid because signaling goes directly from the cellular membrane to the nucleus and path **c** has one invalid interaction because signaling goes in a direction against the assumed signaling flow.

We acknowledge that the assumptions in this work may not hold for many pathways. For example, some pathways are initiated via nuclear receptors, and would be missed based on our assumption that signaling begins at receptors at the cell membrane. We also do not consider other compartments beyond *ExtMem*, *Cytosol*, and *Nucleus* in our model, while the mitochondria and secretory vesicles play an important role in some signaling pathways. These decisions can be taken by the user, which makes the proposed model of signaling flow customizable to a pathway under study. *A priori* information about the structure of signaling flow may further improve *LocPL* predictions.

### 2.4 Dynamic Program for Path-Based Signaling Scores

Given a path *P* = (*v*_1_, *v*_2_*, …, v*_*m*_) that connects *m* proteins, our goal is to find a selection of compartments that maximize the path signaling score (by sum of log-transformed localization scores) while respecting the assumed signaling flow structure outlined above. For each protein *v ∈ V*, we use 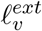, 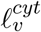 and 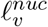 to denote the ComPPI scores of *ExtMem*, *Cytosol*, and *Nucleus* respectively. We log-transform these scores to be localization costs, that is, 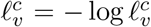 for each protein *v* and each cellular compartment *c* (either *ExtMem*, *Cytosol*, or *Nucleus*). Let *s*(*v*_*j*_, *c*) be the optimal score of the path up to node *v_j_ ∈ P*, where *v*_*j*_ is in compartment *c*. The optimal signaling score of the path must end in the nucleus, which we denote by *s*(*v*_*m*_, *nuc*). Since our assumed signaling model requires that signaling advances through pairs of interacting proteins sharing a cellular compartment or through proteins that traverse multiple compartments, there are only three routes for the signaling information to advance from protein *v*_*m*−1_ to end up in the nucleus for protein *v*_*m*_: 1) protein *v*_*m*−1_ and protein *v*_*m*_ interact in the cytosol and then protein *v*_*m*_ moves to the nucleus, 2) protein *v*_*m*−1_ moves from the cytosol to the nucleus and then interacts with protein *v*_*m*_ in the nucleus, or 3) protein *v*_*m*−1_ and protein *v*_*m*_ interact in the nucleus. Based on these constraints, the optimal path signaling score *s*(*v*_*m*_, *nuc*) can be computed as:

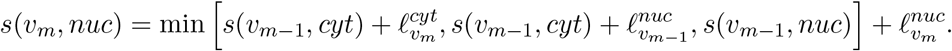

In general, at node *v*_*j*_, *j* = 2, 3, …, (*m* − 1), the set of equations for the scores are:

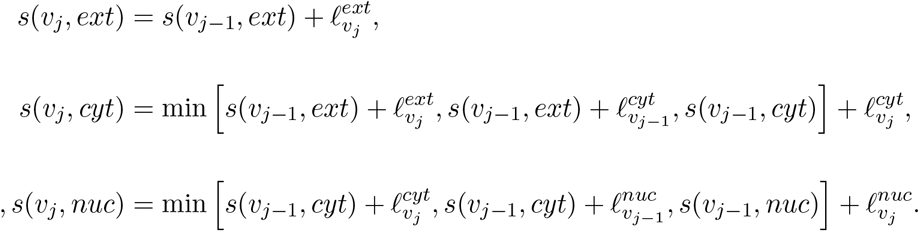

Note that we can only reach a protein in *ExtMem* from another protein in *ExtMem*, we can reach a protein in *Cytosol* from another protein in either *ExtMem* or *Cytosol*, and we can reach a protein in *Nucleus* from another one in either *Cytosol* or *Nucleus*.

To ensure that the path starts with the cellular compartment *ExtMem*, the base case for these recurrence relations are:

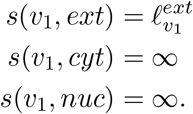

The final score taken will be *s*(*v*_*m*_, *nuc*) since we require the path to terminate in the nucleus. These recurrence relations can be calculated using a dynamic program in linear time w.r.t. the path length for each tied path. An illustrative example of this dynamic program is provided in Supplementary Section 1.

### 2.5 The Color-Coding-Based Method

Color-coding is a randomized technique that computes simple paths that start and end at two different vertices and no vertex is visited more than once [26]. Given a graph *G*, a set *R* of a path starting points (e.g. cellular membrane receptors) and a set *T* of ending points (e.g. transcriptional regulators (TRs)), and a fixed number *l* representing the path length (number of vertices), the color-coding method randomly assigns to each vertex in the graph a uniformly distributed color (label) from {1, 2, …, *l*}, and then finds a *colorful* path that starts at a receptor (*v*_1_ *∈ R*), ends at a TR (*v_l_ ∈ T*), and each one of the *l* vertices composing the path has a distinct color. The constraint of a colorful path (distinct colors of the path vertices) ensures that the reconstructed path is simple. The random designation of colors to the vertices leads to an optimal/sub-optimal solution, if one exists. So, a large number of iterations is required to increase the probability of finding a colorful path. The number of iterations increases exponentially with increasing the probability of success and/or the path length [26]. Enhanced versions of the original color-coding method were proposed to speed up the technique as in [29–31].

The method described in [25] extends the original color-coding technique [26] by integrating proteins cellular information at reconstructing signaling pathways. To the best of our knowledge, that extended color-coding version [25] (called CC from here on) is the closest in its aim to what we propose in this study. Beside the constraint of a colorful path, CC allows signaling to advance across the different cellular compartments in a predefined order, i.e. from the cell membrane to the cytosol and then into the nucleus.

*LocPL* produces *k* paths: the *k*-shortest paths. In order to compare *LocPL* against CC, we need CC to produce the same number of paths, where *k* = 20, 000 in this study. This in turn requires running CC a number of iterations much larger than *k* to account for the trials of non-colorful paths. This can take up to days, if not weeks, for a single pathway when the interactions network is very large. The sped up versions of CC mentioned above were tested against relatively smaller networks with hundreds or a few thousands of edges, and many of them may need much modification to integrate the proteins cellular information. So, we augment CC with Yen’s algorithm [32] to compute the *k*-shortest paths based on the CC method. We call this the *Yen_CC* method. Once Yen’s algorithm finds a path, it searches for alternative paths that differ from the discovered path in one or more edges. In other words it searches for new partial paths. Hence, in *Yen_CC*, instead of running a new iteration to find a complete colorful path, the iteration will look for a partial colorful path, leading to reduction in the search space and time. *Yen_CC* does not handle tied reconstructions, and it reports paths with the same reconstruction cost in an arbitrary order in the *k*-paths list. Details about how we implemented the CC method and how we augmented it with Yen’s algorithm are provided in the Supplementary Section 4.

### 2.6 Interactomes and Pathways

#### *PLNet_2_* Interactome

We built *PLNet*_2_ from both physical molecular interaction data (BioGrid, DIP, InnateDB, IntAct, MINT, PhosphositePlus) and annotated signaling pathway databases (KEGG, NetPath, and SPIKE) [33–37]. *PLNet*_2_ contains 17,168 nodes, 40,016 directed regulatory interactions, and 286,250 bidirected physical interactions, totaling 612,516 directed edges. We assigned interaction direction based on evidence of a directed enzymatic reaction (e.g., phosphorylation, dephosphorylation, ubiquitination) from any of the source databases. Each inter-action is supported by one or more types of experimental evidence (e.g. yeast two hybrid or co-immunoprecipitation), and/or the name of the pathway database. Edges are weighted using an evidence-based Bayesian approach that assigns higher confidence to an experiment type database if it identifies interacting proteins that participate in the same biological process [9]. Given a set *P* of positive edges and a set *N* of negative edges, the method estimates, for each evidence type *t*, the probability that *t* supports positive interactions. These probabilities are then combined for each interaction supported by (potentially multiple) evidence types to produce a final weight. We chose the GO term “regulation of signal transduction” (GO:0009966) to build a set of positive interactions that are likely related to signaling. Positives are edges whose nodes are both annotated with this term, and negatives are randomly selected edges whose nodes are not co-annotated to the term. We chose |*N*| = 10 × |*P*| negative edges. To lessen the influence of very highly-weighted edges, we apply a ceiling of 0.75 to all weights [9].

#### HIPPIE Interactome

HIPPIE (Human Integrated Protein Protein Interaction rEference) is a repository of 16,707 proteins and 315,484 PPIs [2] (version 2.1, July 18^*th*^, 2017 [38]). Each interaction has a confidence score calculated as a weighted sum of the number of studies detecting the interaction, the number and quality of experimental techniques used in these studies to measure the interaction, and the number of non-human organisms in which the interaction was reproduced [2]. We ensure that all NetPath interactions are in HIPPIE by using a tool that is provided on the HIPPIE website [38] to integrate new interactions to HIPPIE. We used that tool to score the missed NetPath interactions with the default parameter values used to score the HIPPIE interactions. This lead to adding 792 proteins and 6,379 PPIs to make HIPPIE of 17,499 and 321,863 PPIs in total.

#### Ground Truth Pathways

We consider a set of four diverse pathways from the NetPath database [35] as our ground truth: *α*6*β*4 Integrin, IL2, EGFR1, and Wnt. Receptors and TRs are automatically detected for each of the eight pathways from lists of 2,124 human receptors and 2,286 human TRs compiled from the literature; see [13] for more details. Supplementary Table 1 summarizes the number of interactions, receptors, and TRs per pathway.

### 2.7 Global and Path-Based Assessment

We assess the performance of *LocPL* compared to PathLinker (*PL*) and *Yen_CC* using two methods that evaluate global and local features of the ranked paths.

#### Precision-recall (PR) curves

Given a ranked list of paths, we order each interaction by the index of the path in which it first appears. We compute precision and recall for this ranked list using the NetPath interactions as positives and a sampled set of negative interactions that are 50 times the size of the positive set.

#### Path-based assessment

The PR curves provide a global quantitative assessment across all the *k* paths in a reconstruction, showing how quickly (in terms of *k*) the technique can discover new positive edges. However, this approach considers a positive only once, i.e., the first times it appears in a path. Thus, this global measure fails to characterize each path individually in terms of the number of positives contained in that path. Hence, we introduce a simple way to “locally” assess paths by computing the within-path percentage of true positive edges, denoted as *PosFrac*. Since we compute this metric value independently for each path, it does not matter if a positive interaction is detected earlier in another path. We compute the *PosFrac* value over non-overlapping windows of paths. For example, for a window of 100 paths, we compute the average *PosFrac* over the first 100 paths, then the average *PosFrac* over the second 100 paths, and so on, providing *k/*100 values to plot.

#### Statistical significance

The global assessment is based on two concurrent values: precision and recall. These two quantities are related, so we use their harmonic mean (*F*_1_ score) to get a single value summarizing both values:

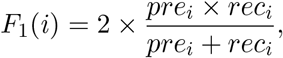

where *pre*_*i*_ and *rec*_*i*_ are the *i*-th values of precision and recall, respectively. The *F*_1_ score values are fed to the Mann-Whitney U (MWU) statistical test for unpaired samples to estimate whether the difference in results between *LocPL* and *PL*, and between *LocPL* and *Yen_CC* is statistically significant. The inputs to the MWU test for the path-based assessment are the *PosFrac* values. We acknowledge that *PosFrac*, precision and recall are not purely independent between the two methods, so there is some dependence introduced in the MWU tests.

## 3 Results

### 3.1 Combining Interactomes with Localization Information

Approximately 95% of the proteins in *PLNet*_2_ have localization information, producing an interactome with about 86% of the edges (Table 1). Only 65% of the HIPPIE proteins have localization information, making a much smaller interactome with only about 34% of the original edges. All pathway receptors and TRs in *PLNet*_2_ have localization information, and nearly all of them (82 out of 91) in HIPPIE have this information (Supplementary Table 1). After filtering *PLNet*_2_ using ComPPI, 62% of the proteins have a non-zero *ExtMem* localization score, 78% have a non-zero *Cytosol* localization score, and 64% have a non-zero *Nucleus* localization score (Supplementary Table 2). Most of the proteins have non-zero localization scores for multiple compartments, though 62% of the proteins with a single non-zero localization score appear in the *Nucleus*.

**Table 1:** Number of proteins and interactions in *PLNet*_2_ and HIPPIE.

| Interactome | Complete Interactome | | Interactome ∩ ComPPI | |
| --- | --- | --- | --- | --- |
|  | Nodes | Edges | Nodes | Edges |
| *PLNet<sub>2</sub>* | 17,168 | 612,516 | 16,225 | 527,706 |
| HIPPIE | 17,499 | 321,863 | 11,430 | 108,391 |

Applying PathLinker to the ComPPI-filtered interactome partially mitigates the problem of tied paths, but many ties remain. For example, after running PathLinker on the *α*6*β*4 Integrin pathway with the full *PLNet*_2_ interactome, there were 82 groups of paths where each group shared the same reconstruction score (Supplementary Table 3). This number was reduced to 58 groups when running PathLinker on the filtered *PLNet*_2_ interactome. However, ties still dominate the reconstruction scores; thus the need for an approach to breaking these ties and re-prioritizing paths in a biologically relevant way is still imperative.

### 3.2 Assessment of Pathway Reconstructions

We applied PathLinker (*PL*) and *LocPL* to signaling pathways from the NetPath database to the *PLNet*_2_ and HIPPIE interactomes as described in Section 2.6 (Interactomes and Pathways). We computed *k* = 20, 000 paths for each approach, similar to the original publication [13]. Paths that have the same reconstruction score differ substantially in their signaling scores computed by the dynamic program. Figure 3 shows four examples of the signaling score *s*_*i*_ distribution for paths with the same reconstruction score *r*_*i*_. Signaling scores are used to re-order paths sharing the same reconstruction score. We also computed 20,000 paths using the *Yen_CC* approach for the *PLNet*_2_ interactome only due to the very long time needed to run *Yen_CC*. We show results for the *PLNet*_2_ interactome first and then show those for HIPPIE.

**Figure 3:**
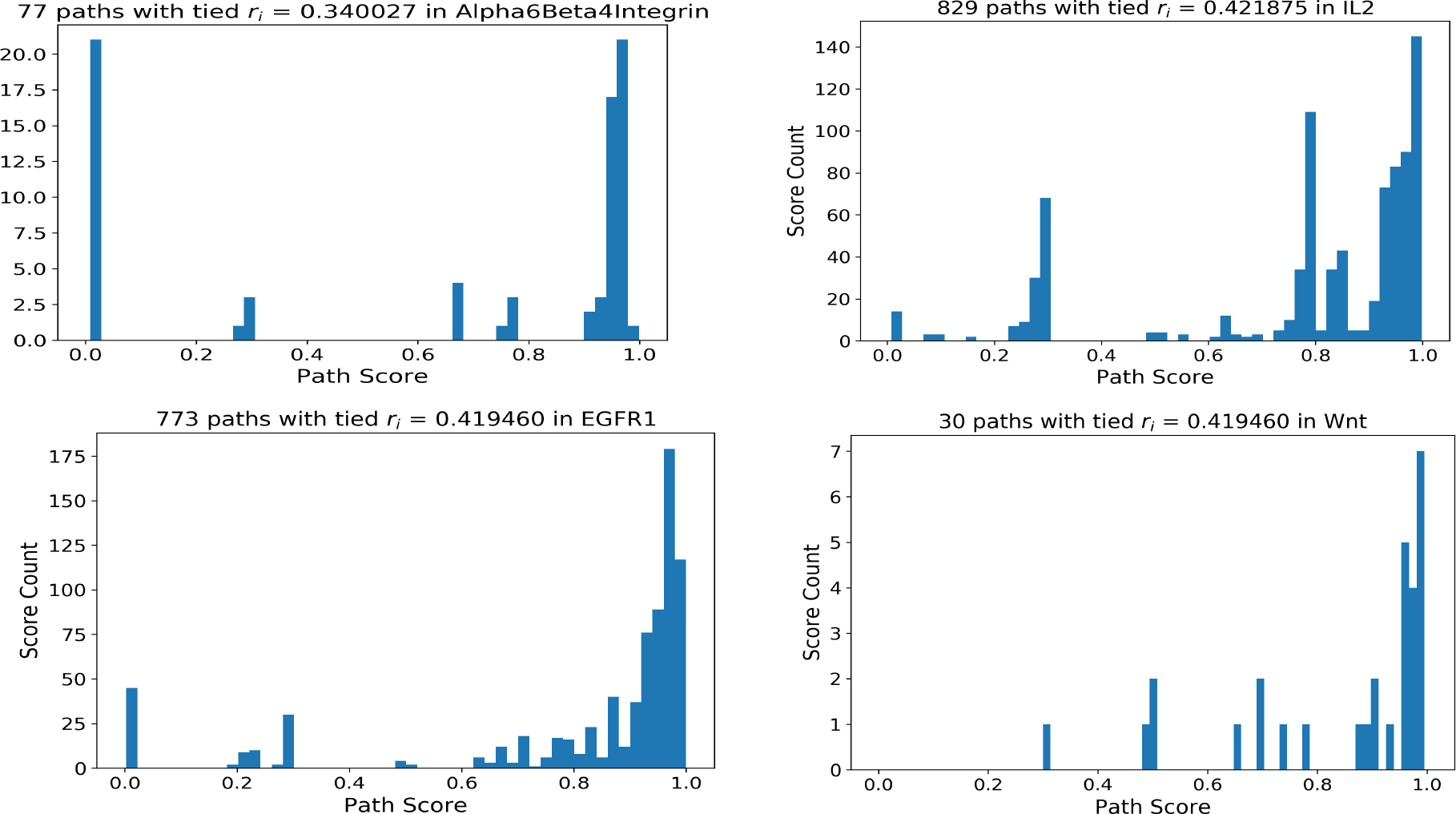
Histogram of signaling scores *s*_*i*_ for paths with tied reconstruction score *r*_*i*_. The titles indicate the pathway name, the *r*_*i*_ value, and the number of paths tied with this *r*_*i*_.

#### Precision and Recall

We assessed *PL*, *LocPL*, and *Yen_CC* using the *PLNet*_2_ interactome on four signaling pathways: *α*6*β*4 Integrin, EGFR1, IL2, and Wnt. *LocPL* generally outperforms *PL* and *Yen_CC* across all four pathways in terms of precision and recall, where the precision of *LocPL* is greater than *PL* and *Yen_CC* at nearly all values of recall (Figure 4 (Left)). Moreover, *LocPL* usually detects higher proportions of positives than *PL* and *Yen_CC* as reflected in the larger recall values for *LocPL* (Figure 4 (Left)), though the same number of paths were recovered for each method. For every value of precision and recall, we plotted the harmonic mean (*F*_1_ score) of the two values in Figure 4 (Right). The *F*_1_ curve for *LocPL* is significantly higher than that of *PL* and *Yen_CC* for the four pathways (MWU test *p*-value ≤ 0.0001).

**Figure 4:**
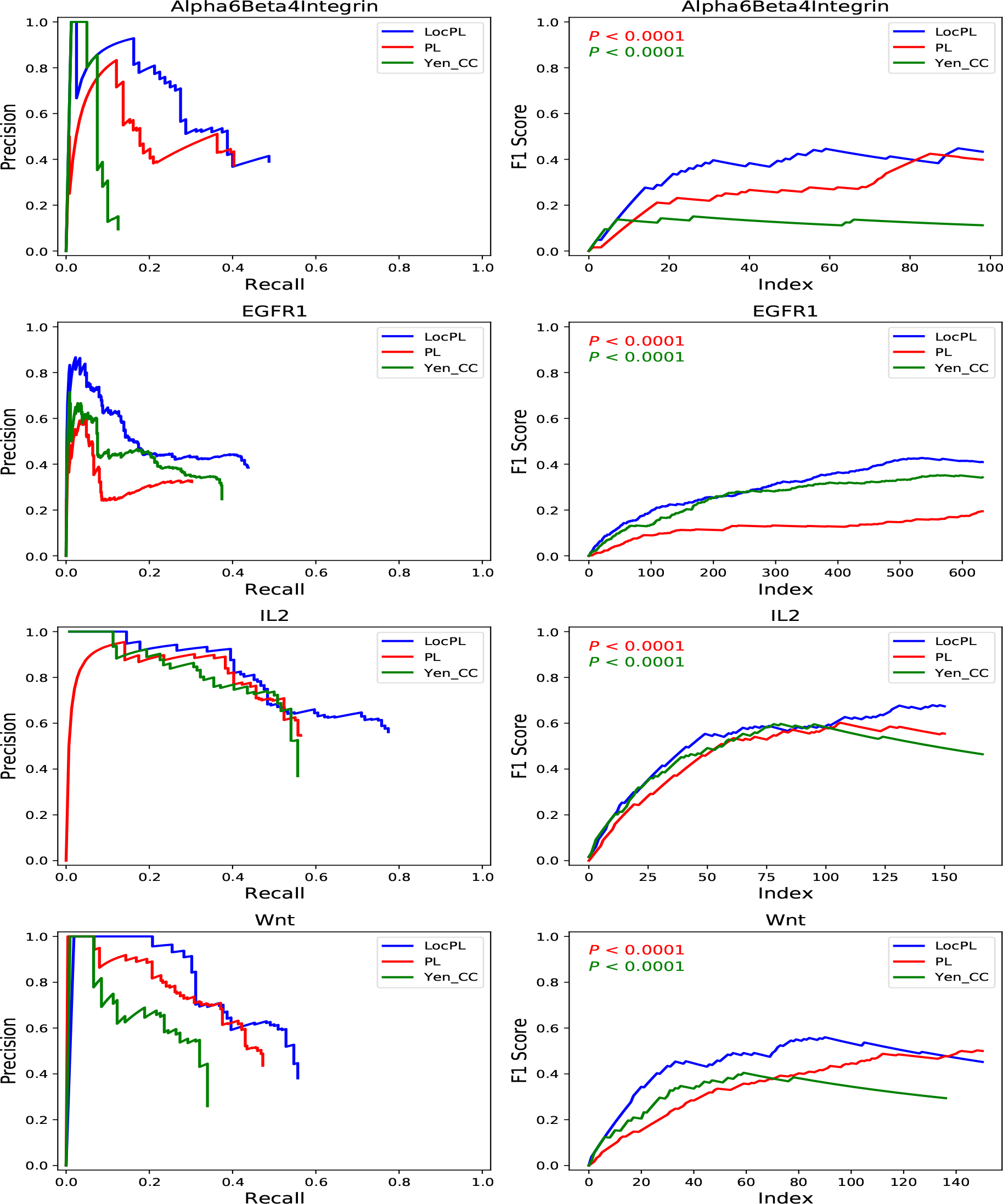
*PLNet*_2_: (**Left**) Precision and recall curves of pathway reconstructions from PathLinker (*PL*), *LocPL*, and *Yen_CC* on four NetPath signaling pathways. (**Right**) *F*_1_ scores for the individual NetPath pathways. These values are fed to the MWU test to check for difference significance. The *p*-value, *P*, is for the MWU test (alternative: *LocPL* > *PL* or *LocPL* > *Yen_CC*). The color of the *p*-value text indicates which method is tested against *LocPL*, e.g. the red text tests that the *F*_1_ score of *LocPL* is greater than that of *PL*.

#### Assessment of Aggregate Pathways

To assess overall effect of *LocPL* on signaling pathway reconstructions, we considered precision and recall aggregated over the four NetPath signaling pathways (Supplementary Section 3) for *PLNet*_2_ (Figure 5 (left)). *LocPL* shows better performance over *PL* and *Yen_CC* at nearly all the *k* values used to compute precision and recall. This improvement is striking at almost all values of recall, with gains in precision that range from 6% to 32% at recall of 0.37 and 0.17, respectively, against *PL*. When compared to *Yen_CC*, *LocPL* achieves gain in precision of about 27% for recall of 0.1 and on. Superiority of *LocPL* is significant (MWU test, Figure 5 (Right)), where the aggregate *F*_1_ score values are higher everywhere for *LocPL*.

**Figure 5:**
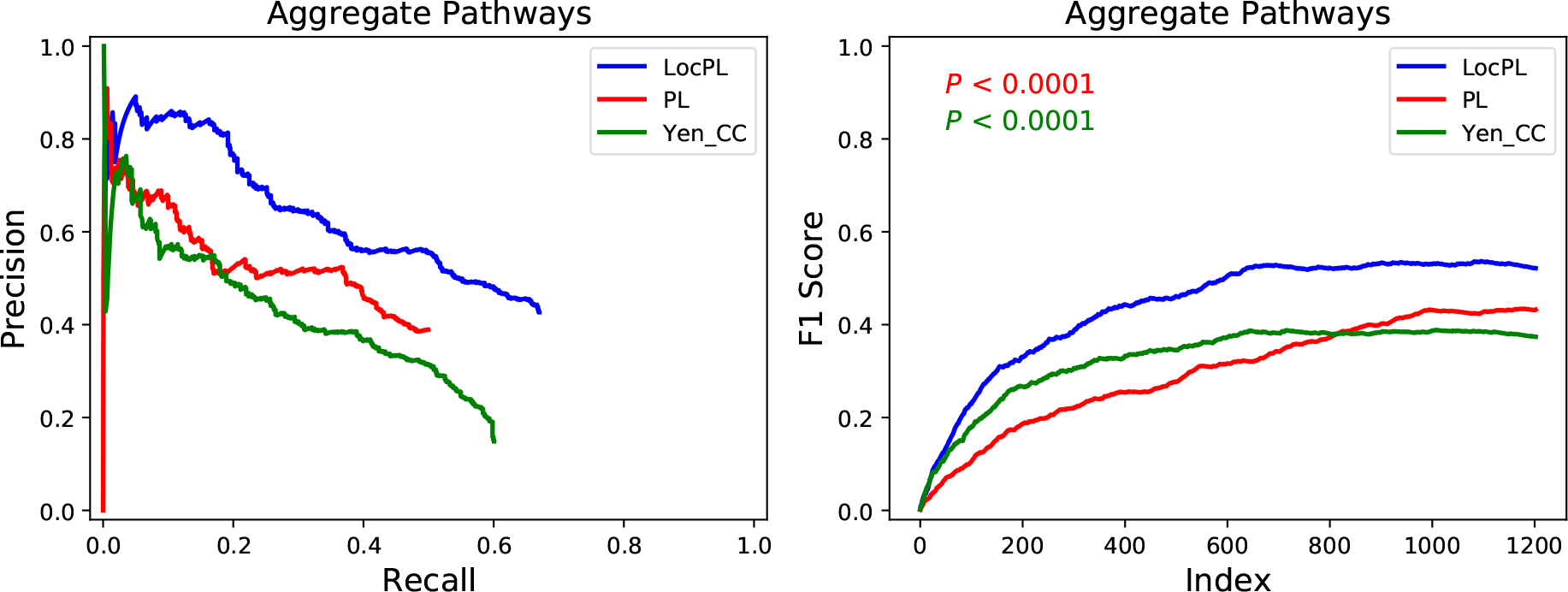
*PLNet*_2_: (**Left**) Precision-Recall curve and (**Right**) *F*_1_ score curve of *PL*, *LocPL*, and *Yen_CC* computed on paths aggregated across all four signaling pathways. The *p*-value, *P*, is for the MWU test (alternative: *LocPL > PL* or *LocPL > Yen CC*). The color of the *p*-value text indicates which method is tested against *LocPL*, e.g. the red text tests that the *F*_1_ score of *LocPL* is greater than that of *PL*.

#### Path-based Assessment

In addition to the global assessment, we are interested in the quality of subsets of paths. Plotting *PosFrac* of non-overlapping windows of 100 paths reveals subsets of paths that are enriched for positive interactions in the four pathway reconstructions (Figure 6).^1^ For example, about more than 80% and 85% of the paths produced by *LocPL* for the IL2 pathway reconstruction tend to contain more positive signaling edges than those obtained by *PL* and *Yen_CC*, respectively, over all the 20,000 paths. *PosFrac* is almost consistent for *LocPL* and, despite some spikes (of different widths) for *PL* and *Yen_CC*, *PosFrac* for *LocPL* dominates the graph (mean *±* standard deviation values of *PosFrac* are 0.23 *±* 0.06, 0.11 *±* 0.12, and 0.14 *±* 0.07 for *LocPL*, *PL*, and *Yen_CC*; respectively). In the IL2 pathway reconstruction, this distinction is significant (one-tailed MWU test, Figure 6). *LocPL* is also significantly better than *PL* and *Yen_CC* for the *α*6*β*4 Integrin and EGFR1 pathways. The situation is different for the Wnt pathway, where *LocPL* is statistically significant when compared against *Yen_CC* (Figure 6 (lower right)), but statistically insignificant when tested against *PL* (*p*-values of 0.9726, Figure 6 (lower left)).

**Figure 6:**
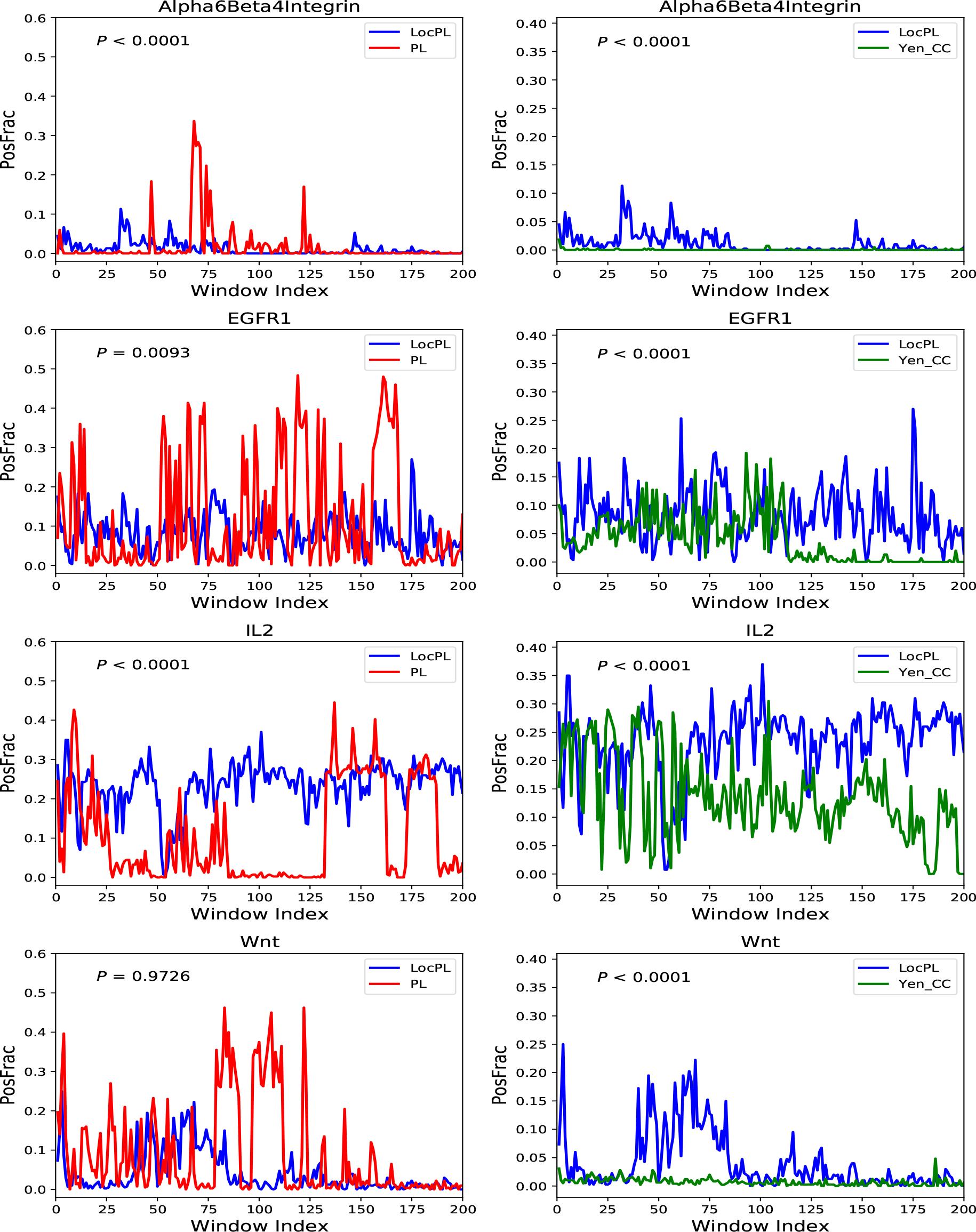
*PLNet*_2_: Path-based performance of four NetPath signaling pathways for (**Left**) *LocPL vs. PL* and (**Right**) *LocPL vs. Yen CC*. *PosFrac* is the percentage of positives averaged across non-overlapping windows of 100 paths. The *p*-value, *P*, is for the MWU test (alternative: *LocPL* > *PL* or *LocPL > Yen CC*).

#### Results on the HIPPIE Interactome

We extended our experiments on the four NetPath signaling pathways (*α*6*β*4 Integrin, EGFR1, IL2, and Wnt) to the HIPPIE interactome. Figure 7A (Left) shows, for all the four pathways, that the precision of *LocPL* is greater than that for *PL*, and that the proportions of positives detected by *LocPL* is always higher than those of *PL*. This consistently leading performance of *LocPL* over *PL* is evidently statistically significant (Figure 7A (Right)). Again, the aggregate precision of *LocPL* has gains of up to 40% over that of *PL*, and the recall proportion is more than the double for *LocPL* (Figure 7C). The reconstructed paths of *LocPL* are steadily and significantly more enriched with positive interactions than the paths of *PL* (Figure 7B).

**Figure 7:**
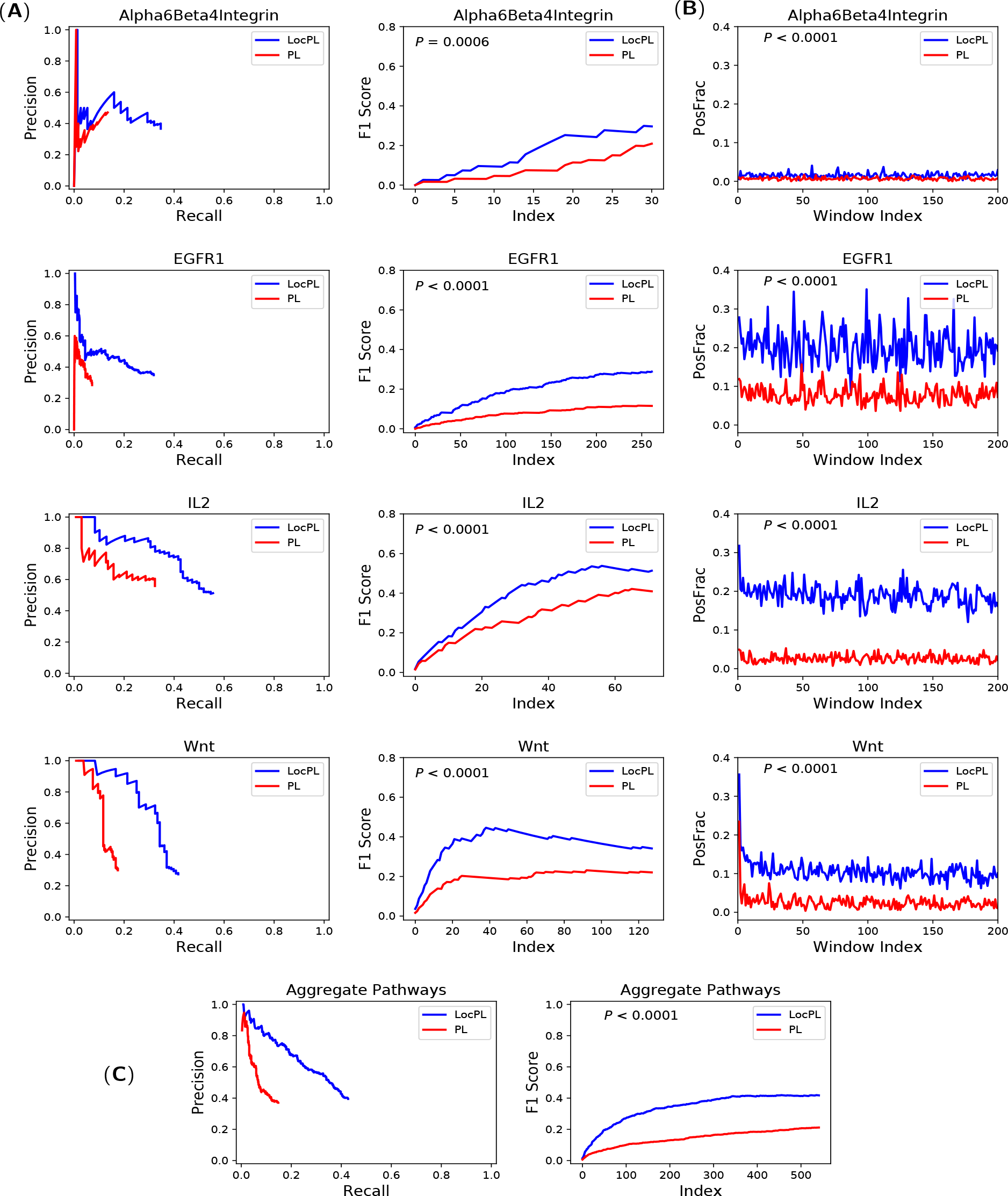
HIPPIE: (**A: Left**) Precision and recall curves of pathway reconstructions from Path-Linker (*PL*) and *LocPL* on four NetPath signaling pathways. (**A: Right**) *F*_1_ scores for the individual NetPath pathways. (**B**) Path-based performance of the individual pathways. *PosFrac* is the percentage of positives averaged across non-overlapping windows of 100 paths. (**C: Left**) Aggregate PR curve, and (**C: Right**) *F*_1_ score curve over the four signaling pathways. The *p*-value, *P*, is for the MWU test (alternative: *LocPL > PL*).

### 3.3 Comparison of Pathway Reconstructions

*LocPL* provides a compartment-aware ranking of paths connecting receptors to TRs. In addition to the global and local assessments provided above, we examined the 100 top-ranking paths of *PL*, *LocPL*, and *Yen_CC* pathway reconstructions using *PLNet*_2_ for the *α*6*β*4 Integrin, IL-2, EGFR1, and Wnt pathways. We first counted the number of paths with at least one positive interaction and the number of paths whose all interactions are positives within the first 10 and 100 paths. In most of the cases, *LocPL* identifies more positive-enriched paths than *PL* and *Yen_CC* (Table 2). Note that the number of positives in the earliest paths for the Wnt pathway is larger for *PL* over *LocPL*, which agrees with the *PosFrac* values shown in Figure 6 (lower left).

**Table 2:**
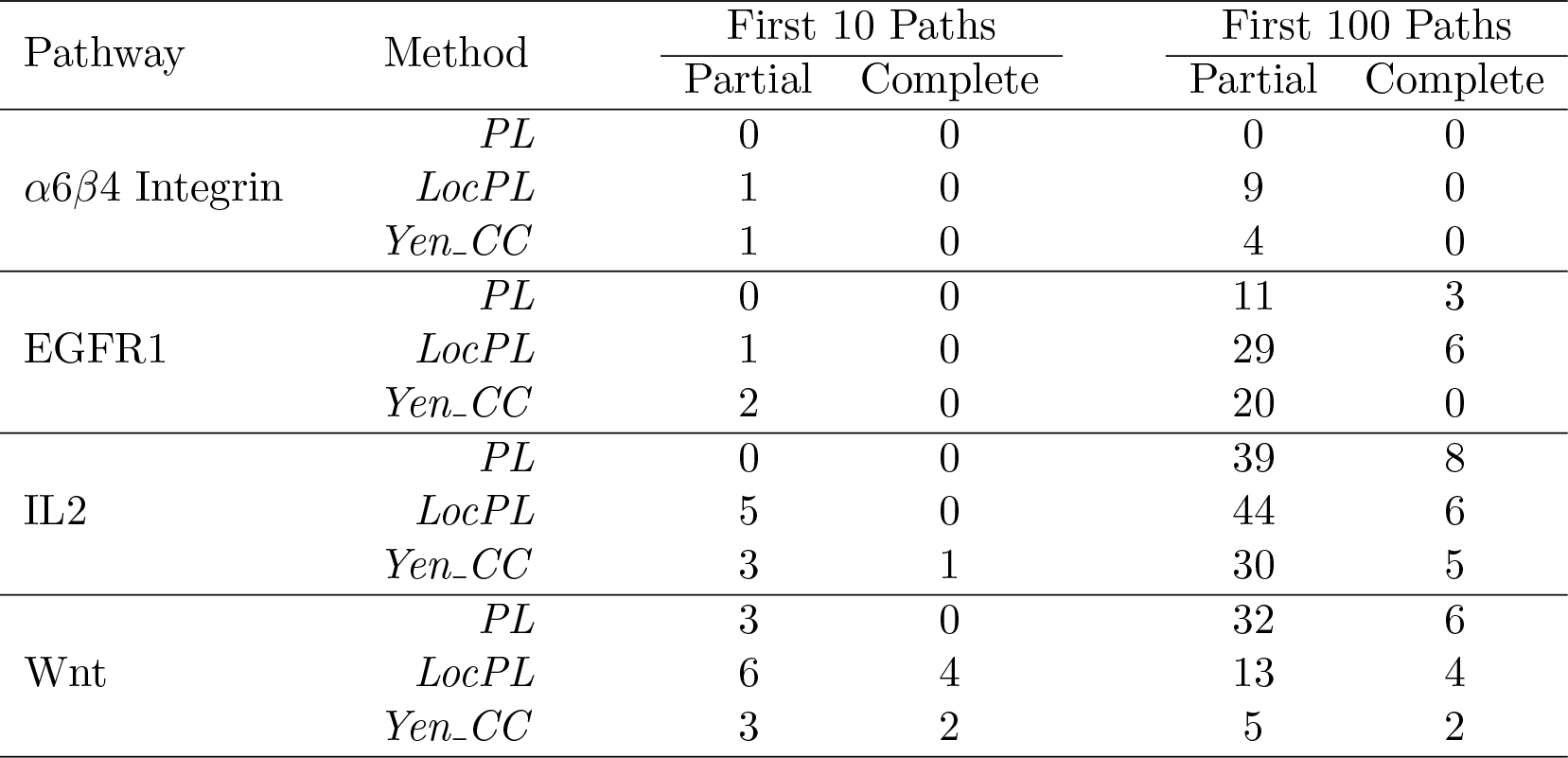
*PLNet*_2_: The number of paths with at least one positive interaction (partial) and with all interactions are positives (complete) among the first 10 and 100 reconstructed paths.

| Pathway | Method | First 10 Paths |  | First 100 Paths |  |
| --- | --- | --- | --- | --- | --- |
|  |  | Partial | Complete | Partial | Complete |
| $\alpha6\beta4$ Integrin | *PL* | 0 | 0 | 0 | 0 |
|  | *LocPL* | 1 | 0 | 9 | 0 |
|  | *Yen_CC* | 1 | 0 | 4 | 0 |
| EGFR1 | *PL* | 0 | 0 | 11 | 3 |
|  | *LocPL* | 1 | 0 | 29 | 6 |
|  | *Yen_CC* | 2 | 0 | 20 | 0 |
| IL2 | *PL* | 0 | 0 | 39 | 8 |
|  | *LocPL* | 5 | 0 | 44 | 6 |
|  | *Yen_CC* | 3 | 1 | 30 | 5 |
| Wnt | *PL* | 3 | 0 | 32 | 6 |
|  | *LocPL* | 6 | 4 | 13 | 4 |
|  | *Yen_CC* | 3 | 2 | 5 | 2 |

We then wished to better understand how the constraints imposed by the dynamic program affected the pathway reconstructions. We compared the subgraph comprised of the first 100 paths before applying the dynamic program that reorders ties based on signaling score, to the subgraph comprised of the first 100 paths after applying the dynamic program. While the number of nodes and edges were about the same between the two subgraphs, we found that EGFR1, IL2, and Wnt only had about half the number of nodes in common and about a third the number of edges in common (Supplementary Figure 2). The number of common nodes and edges for the two subgraphs of *α*6*β*4 Integrin are about, at least, double the number of the unique nodes and edges to either subgraph.

We also visualized networks for each pathway reconstruction before and after applying the dynamic program (Figure 8). The nodes are colored according to red, green, and blue channels depending on the ComPPI localization scores for membrane, cytosol, and nucleus respectively; a protein that appears in all compartments will be white. The signaling flow constraints from the dynamic program on *LocPL* paths imply two features about these networks: first, the node colors should change from red (membrane) to green (cytosol) to blue (nucleus), and second, no paths of length one are allowed. Both of these features are visible in the comparison of the IL2 pathway reconstructions (Figure 8A). For example, the edge from IL2 Receptor A (IL2RA) to transcription factor STAT5B is removed after the dynamic program, removing the IL2RA receptor from the first 100 paths.

**Figure 8:**
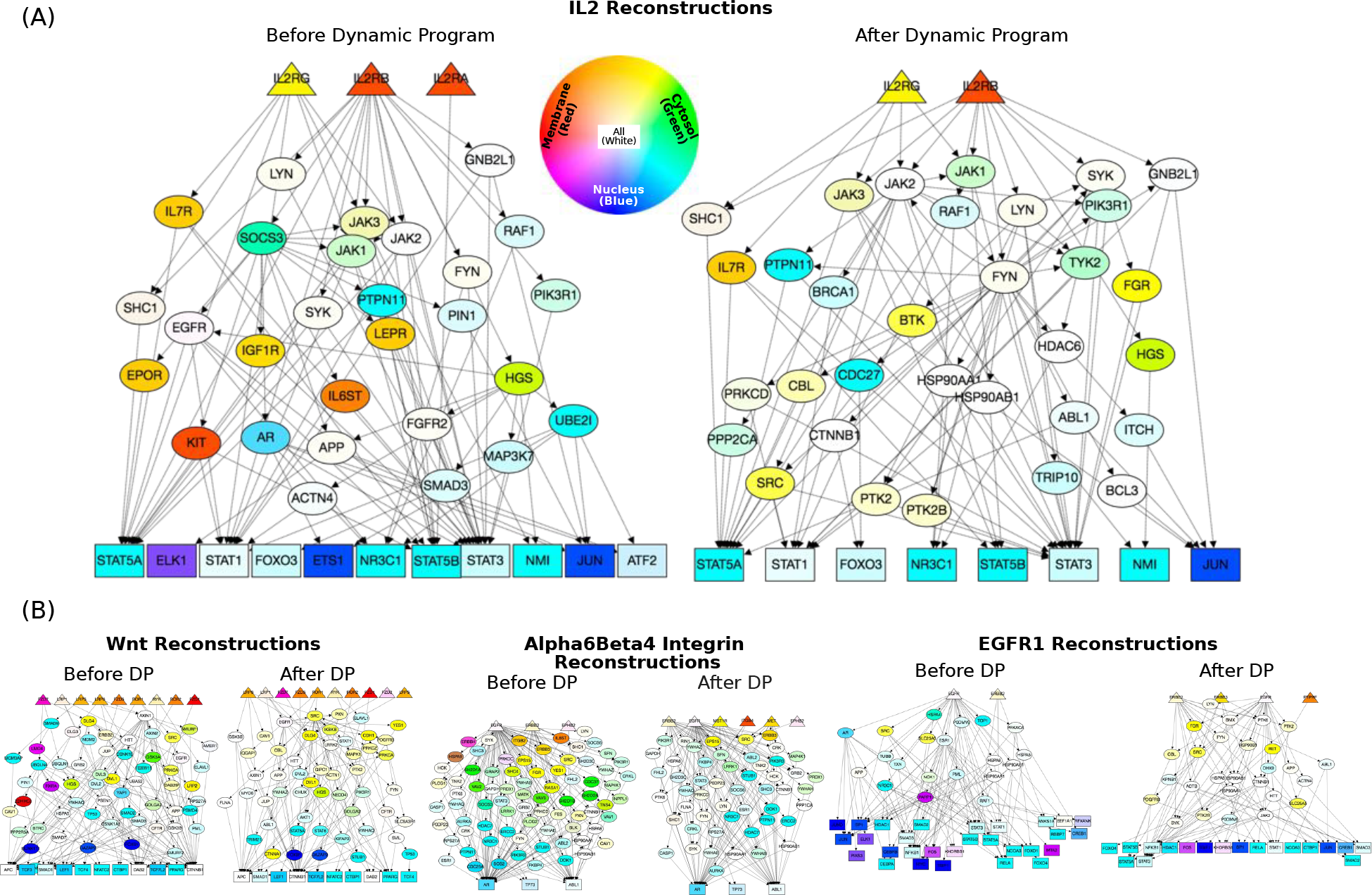
*PLNet*_2_: *LocPL* pathway reconstructions (first 100 paths). **(A)** IL2 pathway reconstructions before applying the dynamic program (left) compared to after applying the dynamic program (right). **(B)** Topologies of other pathway reconstructions; larger figures provided in Supplementary Figures 3-5. Receptors are labeled as triangles, transcriptional regulators are rectangles, intermediary proteins are ellipses. Color denotes compartment localization; proteins may belong to multiple compartments (and will be lighter shades). Networks were generated using GraphSpace [39], and are available at http://graphspace.org/graphs/?query=tags:LocPL.

The color differences between the two IL2 networks are also notable. Before the dynamic program, the IL2 reconstruction contains main proteins that are predicted to be at the membrane, including the IL7 receptor (IL7R), Insulin Like Growth Factor 1 Receptor (IGF1R), Leptin Receptor (LEPR), KIT Proto-Oncogene Receptor Tyrosine Kinase (KIT), and Erythropoietin Receptor (EPOR). Further, the Interleukin 6 Signal Transducer (IL6ST) is also reported to be at the membrane, yet is downstream of Suppressor Of Cytokine Signaling 3 (SOCS3) in the network (Figure 8A (Left)). IL2 signaling activates the Jak/STAT pathway, and many paths containing Janus kinase family members (JAK1, JAK2, JAK3) also include SOCS3 upstream of these proteins. After the paths are reordered according to the dynamic program, the JAK proteins are directly dosntream of the receptors (Figure 8A (Right)). While some receptors remain after reordering, they either directly interact with the IL2 receptors (e.g. IL7R), or they lie downstream of a protein that is consistent in terms of the signaling constraints. For example, the the SYK-FGR is allowable because SYK has a large ComPPI score for all compartments. The other pathways exhibit dramatic differences in topology compared to the IL2 reconstructions, including the large number of receptors in the Wnt reconstructions, the large number of TFs in the EGFR1 reconstructions, and the large number of intermediate nodes in the Alpha6 *β*4 Integrin reconstruction (Figure 8B and Supplementary Figures 3, 4, and 5).

## 4 Discussion

We present *LocPL*, an automatic signaling reconstruction algorithm that incorporates information about protein localization within the cell. Previous reconstructions contained many tied paths. *LocPL* overcomes this obstacle with a computational framework that favors paths that follow specific assumptions of signaling flow. This framework includes filtering interactions based on their predicted interaction score and applying a dynamic program to each path that finds the most likely series of cellular compartments that are consistent with the model of signaling flow.

Using a new interactome, *PLNet*_2_, we have shown that *LocPL* pathway reconstructions for four pathways are more enriched with positive interactions than paths computed by *PL* and by a peer method, *Yen_CC*, based on the color coding technique. Precision of *LocPL* dominates the precision of *PL* and *Yen_CC* at nearly every value of recall (Figure 4 (Left)), and the resulting *F*_1_ scores are significantly better for *LocPL* (Figure 4 (Right)). *LocPL* dramatically improves precision at all values of recall across four signaling pathways, and this difference is significant by the MWU test (Figure 5).

In addition to the precision and recall assessment used previously by PathLinker [13], we proposed a measure, *PosFrac*, to assess individual paths in terms of proportion of positive signaling interactions. PR curves demonstrate how quickly positive interactions are recovered in a reconstruction, but do not consider the fact that many paths may contain the same positive. *PosFrac* is a path-based measure that considers the proportion of positives within a set of paths, demonstrating that some sets of paths are enriched for positive interactions that may have appeared in a higher-ranked path. *LocPL* paths are consistently enriched with positive interactions more than the paths reconstructed by *Yen_CC* for all the four signaling pathways, and more than the paths of *PL* for two of the pathways (Figure 6). This measure offers complementary insights to the pathway reconstructions beside the PR curves. For example, paths within windows 50 to 65 for the IL2 pathway (Figure 6) have very small *PosFrac* values among all the 20,000 paths. These paths contain interactions that are not labeled as positives but are “close” to the pathway in some sense, suggesting candidate interactions that may point to non-canonical branches of signaling.

Though both *LocPL* and the color coding method (*CC*, [25]) use protein localization information, but the way this information is employed differs substantially. *CC* uses a binarized version of the localization information; what cellular compartments a protein can be found within. This leads to tied reconstructions due to the deprivation from having other measures, beside the reconstruction cost, to re-prioritize ties. In contrast, *LocPL* uses a probabilistic form of the localization information; the likelihood of a protein to be found in one cellular compartment. This furnishes *LocPL* with a second measure, the signaling score, to untangle ties and re-order reconstructions.

*LocPL* ensures that the constituting interactions, from a receptor to a TR, are spatially-coherent within the different cellular compartments. This feature increases the number of paths that contain positives early in the pathway reconstruction, which supports our hypothesis that *LocPL* locally promotes paths with higher proportions of positives up in the *k*-shortest paths list (Table 2).

*LocPL* is not restricted to our proposed interactome, *PLNet*_2_. We applied *LocPL* to the HIPPIE interactome [2]. We compared *LocPL* to only *PL* due to the very long time demand of the *Yen_CC* method. *LocPL*’s performance was statistically significantly better than *PL* as depicted in the PR and the *F*_1_ score curves (Figure 7(A)) and in the *PosFrac* curves (Figure 7(B)) for the individual NetPath signaling pathways. Moreover, this trend is consistent across the four signaling pathways as well (Figure 7(C)).

In this work, we chose to impose an ordering on a subset of the available compartments from ComPPI (*ExtMem*, *Cytosol*, and *Nucleus*). There are many ways to impose a compartmental ordering of signaling flow to capture other features of signaling, including mitochondria-dependent signaling, nuclear receptor signaling and extracellular signaling. *LocPL* is generalizable to different signaling models, as long as the user specifies compartment relationships in a memoryless manner (the signaling score at the next node depends only on the localization score of the next node and the signaling score at the current node; ignoring signaling score history at previous nodes). To illustrate this point, we developed a model of signaling that also includes the mitochondria compartment. We did not notice any changes in the results when we included the mitochondria into our signaling model, most likely due to the relatively few number of proteins in *PLNet*_2_ that had non-zero *Mitochondria* localization scores (Supplementary Table 2). Details about how this modified signaling model and the dynamic program can be found in Supplementary Section 2.

Visual inspection of the subgraphs containing the first 100 paths in the pathway reconstructions before and after applying the dynamic program reveal that reordering tied paths changes the first 100 paths dramatically, even though the number of nodes and edges remain similar (Supplementary Figure 2). In particular, the dynamic program removes membrane-bound receptors that appear downstream of cytosolic proteins, which can be seen by visual inspection (Figure 8). These and other features can be explored in such network reconstructions.

## 5 Conclusion

In this study, we presented *LocPL*, which is a powerful tool for automatic reconstruction of signaling pathways from protein-protein interactions that leverages the proteins cellular localization information. *LocPL* showed profound and significant better reconstructions over those by peer methods in terms of the total number of the true protein interactions across the whole pathway reconstructions and the number of positive interactions per individual paths with a reconstruction. The framework that we have developed may be extended to other graph-theoretic approaches that return subnetworks of directed structure with an associated reconstruction score, such as trees [11, 10, 15]. Our approach encourages the enumeration of many tied results, since incorporating protein compartment information will help break these ties with biologically relevant information. In addition, we anticipate to develop the technique to compare paths in different contexts, such as tissue-specific or disease-specific signaling.

## Acknowledgements

We thank T.M. Murali for useful discussions about PathLinker and other pathway reconstruction methods.

## Availability of data and material

All code and input files, including the newly-released interactome *PLNet*_2_, are available at https://github.com/annaritz/localized-pathlinker.

## Author’s contributions

AR supervised the project. AR and IY conceived of the project and developed the *LocPL* algorithm. JL generated and weighted the new *PLNet*_2_ network. IY implemented the methods and conducted all experiments. All authors contributed to the writing and editing of the manuscript.

1 Note that *PosFrac* considers all negative interactions for each path, unlike the PR curves in Figure 4 that subsample the negative set of interactions. Thus, the *PosFrac* values will be smaller than one would expect based on the PR curves.

