## Supplementary Material for "Integrating Protein Localization with Automated Signaling Pathway Reconstruction"

**for**

#### 1 Illustrative Example of the Dynamic Program

In this section we give a simple, illustrative example of the dynamic program to break tied paths described in the Methods subsection "Dynamic Program for Path-Based Signaling Scores" in the main manuscript. We first summarize the dynamic program as described in the text, then modify it to work with the original localization scores for ease of exposition. We then walk through scoring a four-edge path with this approach.

Consider the set $V = \{v_1, v_2, \ldots\}$ of proteins that contain localization information (e.g. localization scores for at least one of *ExtMem, Cytosol, and Nucleus*). For each protein $v$, we use $\ell_v^{ext}, \ell_v^{cyt}$, and $\ell_v^{nuc}$ to denote these scores, where $0 \leq \ell \leq 1$ for all scores. We log-transform these scores, that is, $\mathcal{T}_v^c = -\log \ell_v^c$ for each protein $v$ and each cellular compartment $c$. Our goal is to find a selection of compartments that maximize the path score by (by summing log-transformed scores) while respecting the signaling flow structure outlined in the Methods subsection "Signaling Flow Structure and Assumptions" in the main manuscript. Let $v_1, v_2, \ldots, v_m$ be the $m$ proteins in path $P_i$.We aim to compute the optimal signaling score of the entire path ending in the nucleus, which we denote by $s(v_m, nuc)$. Since our assumed signaling model requires that signaling advances through pairs of interacting proteins sharing a cellular compartment or through proteins that traverse multiple compartments, there are only three routes for the signaling information to advance from protein $v_{m-1}$ to end up in the nucleus of protein $v_m$: 1) protein $v_{m-1}$ and protein $v_m$ interact in the cytosol and then protein $v_m$ moves to the nucleus, 2) protein $v_{m-1}$ moves from the cytosol to the nucleus and then interacts with protein $v_m$ in the nucleus, or 3) protein $v_{m-1}$ and protein $v_m$ interact in the nucleus. Based on these constraints, the optimal path signaling score $s(v_m, nuc)$ can be computed as:

$$s(v_m, nuc) = \min\left[s(v_{m-1}, cyt) + \mathcal{T}_{v_m}^{cyt}, s(v_{m-1}, cyt) + \mathcal{T}_{v_{m-1}}^{nuc}, s(v_{m-1}, nuc)\right] + \mathcal{T}_{v_m}^{nuc}. \tag{1}$$

In general, at node $v_j$, $j = 2, 3, \ldots, (m-1)$, the set of equations for the scores are:

$$s(v_j, ext) = s(v_{j-1}, ext) + \mathcal{T}_{v_j}^{ext} \tag{2}$$

$$s(v_j, cyt) = \min\left[s(v_{j-1}, ext) + \mathcal{T}_{v_j}^{ext}, s(v_{j-1}, ext) + \mathcal{T}_{v_{j-1}}^{cyt}, s(v_{j-1}, cyt)\right] + \mathcal{T}_{v_j}^{cyt} \tag{3}$$

$$s(v_j, nuc) = \min\left[s(v_{j-1}, cyt) + \mathcal{T}_{v_j}^{cyt}, s(v_{j-1}, cyt) + \mathcal{T}_{v_{j-1}}^{nuc}, s(v_{j-1}, nuc)\right] + \mathcal{T}_{v_j}^{nuc}. \tag{4}$$

Note that we can only reach a protein in *ExtMem* from another protein in *ExtMem*, we can reach a protein in *cytosol* from another protein in either *ExtMem* or *cytosol*, and we can reach a protein in *nucleus* from another one in either *cytosol* or *nucleus*.

To ensure that the path starts with the cellular compartment *ExtMem*, the base case for these recurrence relations are:

$$s(v_1, ext) = \mathcal{T}_{v_1}^{ext} \tag{5}$$

$$s(v_1, cyt) = \infty \tag{6}$$

$$s(v_1, nuc) = \infty. \tag{7}$$

For ease of exposition, we will work with the localization scores in their original format without log-transforming them. In this case, addition in Equations (1–4) will be multiplication and *min* will be *max*. Moreover ($\infty$) in Equations (6–7) will be ($-\infty$). Here are the equations in their new structure.

$$s(v_m, nuc) = \max\left[s(v_{m-1}, cyt) * \ell_{v_m}^{cyt}, s(v_{m-1}, cyt) * \ell_{v_{m-1}}^{nuc}, s(v_{m-1}, nuc)\right] * \ell_{v_m}^{nuc}. \tag{8}$$

$$s(v_j, ext) = s(v_{j-1}, ext) * \ell_{v_j}^{ext} \tag{9}$$

$$s(v_j, cyt) = \max\left[s(v_{j-1}, ext) * \ell_{v_j}^{ext}, s(v_{j-1}, ext) * \ell_{v_{j-1}}^{cyt}, s(v_{j-1}, cyt)\right] * \ell_{v_j}^{cyt} \tag{10}$$

$$s(v_j, nuc) = \max\left[s(v_{j-1}, cyt) * \ell_{v_j}^{cyt}, s(v_{j-1}, cyt) * \ell_{v_{j-1}}^{nuc}, s(v_{j-1}, nuc)\right] * \ell_{v_j}^{nuc}. \tag{11}$$

$$s(v_1, ext) = \ell_{v_1}^{ext} \tag{12}$$

$$s(v_1, cyt) = -\infty \tag{13}$$

$$s(v_1, nuc) = -\infty. \tag{14}$$

These recurrence relations can be efficiently calculated using a dynamic program, filling an $m \times 3$ table denoting the number of nodes ($m$) by the three compartments. The final score taken will be $s(v_m, nuc)$, since we require that the path terminates in the nucleus.

The following is an example of a path of five nodes/proteins, $\langle v_1, v_2, \ldots, v_5\rangle$, and four edges/interactions. The $5 \times 3$ table below represents the table used to iteratively compute the signaling score. Each column represents a protein and each row represents a cellular compartment. The localization scores (probabilities of a protein to be found in each of the three cellular compartments) are shown above the proteins. The red cells in the table represent the cells that do not affect computing the signaling score. For example, at the first protein in the path $v_1$, we ignore the *Cytosol* and the *Nucleus* compartments to force the path to start with a protein at either the extracellular domain or the cell membrane, and hence both cells take extreme values like ($-\infty$).

**At $v_1$:**

To initialize the dynamic program, we apply Equations (12–14) as shown below in the first column of the table.

| | | | | | |
|---|---|---|---|---|---|
| $Pr\{$Ext$\}$: | $\frac{1}{2}$ | $\frac{1}{2}$ | $\frac{1}{4}$ | $\frac{1}{10}$ | $\frac{1}{2}$ |
| $Pr\{$Cyt$\}$: | $\frac{3}{4}$ | $\frac{3}{4}$ | $\frac{1}{2}$ | $\frac{1}{4}$ | $\frac{3}{4}$ |
| $Pr\{$Nuc$\}$: | $\frac{3}{4}$ | $\frac{1}{4}$ | $\frac{1}{4}$ | $\frac{1}{8}$ | $\frac{1}{4}$ |
| | $v_1$ | $v_2$ | $v_3$ | $v_4$ | $v_5$ |
| *ExtMem* | $\frac{1}{2}$ | — | — | — | |
| *Cytosol* | $-\infty$ | — | — | — | |
| *Nucleus* | $-\infty$ | $-\infty$ | — | — | — |

**At $v_2$:**

From $v_2$ to the end of the path, we use Equations (9–11) to compute the signaling score at the intermediate proteins.

$$
\begin{aligned}
s(v_2, ext) &= s(v_1, ext) * \ell_{v_2}^{ext} = \frac{1}{2} * \frac{1}{2} = \frac{1}{4}. \\
s(v_2, cyt) &= \max\left[s(v_1, ext) * \ell_{v_2}^{ext}, s(v_1, ext) * \ell_{v_1}^{cyt}, s(v_1, cyt)\right] * \ell_{v_2}^{cyt} \\
&= max\left[\frac{1}{2} * \frac{1}{2}, \frac{1}{2} * (-\infty), -\infty\right] * \frac{3}{4} = \frac{3}{16} \\
s(v_2, nuc) &= \max\left[s(v_1, cyt) * \ell_{v_2}^{cyt}, s(v_1, cyt) * \ell_{v_1}^{nuc}, s(v_1, nuc)\right] * \ell_{v_2}^{nuc} \\
&= max\left[(-\infty) * \frac{3}{4}, (-\infty) * \frac{3}{4}, -\infty\right] * \frac{1}{4} = -\infty.
\end{aligned}
$$

Tails of the blue arrows in the table indicate the previous step compartment. The arrow heads indicate the compartment at the current step. The compartments beneath the arrow whole body show the signaling flow across the compartments.

| | | | | | |
|---|---|---|---|---|---|
| $Pr\{\text{Ext}\}$: | $\frac{1}{2}$ | $\frac{1}{2}$ | $\frac{1}{4}$ | $\frac{1}{10}$ | $\frac{1}{2}$ |
| $Pr\{\text{Cyt}\}$: | $\frac{3}{4}$ | $\frac{3}{4}$ | $\frac{1}{2}$ | $\frac{1}{4}$ | $\frac{3}{4}$ |
| $Pr\{\text{Nuc}\}$: | $\frac{3}{4}$ | $\frac{1}{4}$ | $\frac{1}{4}$ | $\frac{1}{8}$ | $\frac{1}{4}$ |
| | $v_1$ | $v_2$ | $v_3$ | $v_4$ | $v_5$ |
| *ExtMem* | $\frac{1}{2}$ | $\frac{1}{4}$ | — | — | |
| *Cytosol* | $-\infty$ | $\frac{3}{16}$ | — | — | |
| *Nucleus* | $-\infty$ | $-\infty$ | — | — | — |

**At $v_3$:**

Following the same procedure used at $v_2$, we get the next equations.

$$
\begin{aligned}
s(v_3, ext) &= s(v_2, ext) * \ell_{v_3}^{ext} = \frac{1}{4} * \frac{1}{4} = \frac{1}{16}. \\
s(v_3, cyt) &= \max\left[s(v_2, ext) * \ell_{v_3}^{ext}, s(v_2, ext) * \ell_{v_2}^{cyt}, s(v_2, cyt)\right] * \ell_{v_3}^{cyt} \\
&= max\left[\frac{1}{4} * \frac{1}{4}, \frac{1}{4} * \frac{3}{4}, \frac{3}{16}\right] * \frac{1}{2} = \frac{3}{32} \\
s(v_3, nuc) &= \max\left[s(v_2, cyt) * \ell_{v_3}^{cyt}, s(v_2, cyt) * \ell_{v_2}^{nuc}, s(v_2, nuc)\right] * \ell_{v_3}^{nuc} \\
&= max\left[\frac{3}{16} * \frac{1}{2}, \frac{3}{16} * \frac{1}{4}, -\infty\right] * \frac{1}{4} = \frac{3}{128}.
\end{aligned}
$$

Multiple arrows of the same color means routes of equal cost for the signaling to flow through across the cellular compartments.

| | $v_1$ | $v_2$ | $v_3$ | $v_4$ | $v_5$ |
|---|---|---|---|---|---|
| $Pr\{\text{Ext}\}$: | $\frac{1}{2}$ | $\frac{1}{2}$ | $\frac{1}{4}$ | $\frac{1}{10}$ | $\frac{1}{2}$ |
| $Pr\{\text{Cyt}\}$: | $\frac{3}{4}$ | $\frac{3}{4}$ | $\frac{1}{2}$ | $\frac{1}{4}$ | $\frac{3}{4}$ |
| $Pr\{\text{Nuc}\}$: | $\frac{3}{4}$ | $\frac{1}{4}$ | $\frac{1}{4}$ | $\frac{1}{8}$ | $\frac{1}{4}$ |
| *ExtMem* | $\frac{1}{2}$ | $\frac{1}{4}$ | $\frac{1}{16}$ | — | |
| *Cytosol* | $-\infty$ | $\frac{3}{16}$ | $\frac{3}{32}$ | — | |
| *Nucleus* | $-\infty$ | $-\infty$ | $\frac{3}{128}$ | — | — |

**At $v_4$:**

$$s(v_4, ext) = s(v_3, ext) * \ell_{v_4}^{ext} = \frac{1}{16} * \frac{1}{10} = \frac{1}{160}.$$

$$s(v_4, cyt) = \max\left[s(v_3, ext) * \ell_{v_4}^{ext}, s(v_3, ext) * \ell_{v_3}^{cyt}, s(v_3, cyt)\right] * \ell_{v_4}^{cyt}$$

$$= max\left[\frac{1}{16} * \frac{1}{10}, \frac{1}{16} * \frac{1}{2}, \frac{3}{32}\right] * \frac{1}{4} = \frac{3}{128}$$

$$s(v_4, nuc) = \max\left[s(v_3, cyt) * \ell_{v_4}^{cyt}, s(v_3, cyt) * \ell_{v_3}^{nuc}, s(v_3, nuc)\right] * \ell_{v_4}^{nuc}$$

$$= max\left[\frac{3}{32} * \frac{1}{4}, \frac{3}{32} * \frac{1}{4}, \frac{3}{128}\right] * \frac{1}{8} = \frac{3}{1024}.$$

| | $v_1$ | $v_2$ | $v_3$ | $v_4$ | $v_5$ |
|---|---|---|---|---|---|
| $Pr\{\text{Ext}\}$: | $\frac{1}{2}$ | $\frac{1}{2}$ | $\frac{1}{4}$ | $\frac{1}{10}$ | $\frac{1}{2}$ |
| $Pr\{\text{Cyt}\}$: | $\frac{3}{4}$ | $\frac{3}{4}$ | $\frac{1}{2}$ | $\frac{1}{4}$ | $\frac{3}{4}$ |
| $Pr\{\text{Nuc}\}$: | $\frac{3}{4}$ | $\frac{1}{4}$ | $\frac{1}{4}$ | $\frac{1}{8}$ | $\frac{1}{4}$ |
| *ExtMem* | $\frac{1}{2}$ | $\frac{1}{4}$ | $\frac{1}{16}$ | $\frac{1}{160}$ | |
| *Cytosol* | $-\infty$ | $\frac{3}{16}$ | $\frac{3}{32}$ | $\frac{3}{128}$ | |
| *Nucleus* | $-\infty$ | $-\infty$ | $\frac{3}{128}$ | $\frac{3}{1024}$ | — |

### At $v_5$:

This is the last node, so we use Equatin (8).

$$s(v_5, nuc) = \max\left[s(v_4, cyt) * \ell_{v_5}^{cyt}, s(v_4, cyt) * \ell_{v_4}^{nuc}, s(v_4, nuc)\right] * \ell_{v_5}^{nuc}$$
$$= max\left[\frac{3}{128} * \frac{3}{4}, \frac{3}{128} * \frac{1}{8}, \frac{3}{1024}\right] * \frac{1}{4} = \frac{9}{2048}.$$

| | | | | | |
|---|---|---|---|---|---|
| $Pr\{\text{Ext}\}$: | $\frac{1}{2}$ | $\frac{1}{2}$ | $\frac{1}{4}$ | $\frac{1}{10}$ | $\frac{1}{2}$ |
| $Pr\{\text{Cyt}\}$: | $\frac{3}{4}$ | $\frac{3}{4}$ | $\frac{1}{2}$ | $\frac{1}{4}$ | $\frac{3}{4}$ |
| $Pr\{\text{Nuc}\}$: | $\frac{3}{4}$ | $\frac{1}{4}$ | $\frac{1}{4}$ | $\frac{1}{8}$ | $\frac{1}{4}$ |
| | $v_1$ | $v_2$ | $v_3$ | $v_4$ | $v_5$ |
| *ExtMem* | $\frac{1}{2}$ | $\frac{1}{4}$ | $\frac{1}{16}$ | $\frac{1}{160}$ | $\frac{1}{320}$ |
| *Cytosol* | $-\infty$ | $\frac{3}{16}$ | $\frac{3}{32}$ | $\frac{3}{128}$ | $\frac{9}{512}$ |
| *Nucleus* | $-\infty$ | $-\infty$ | $\frac{3}{128}$ | $\frac{3}{1024}$ | $\frac{9}{2048}$ |

##### Recovering the Most Probable Compartments:

We can now trace back the most probable route for the signaling flow across the different cellular compartments as shown below by the blue arrows. The signaling score for this path is $\frac{9}{2048}$

| | | | | | |
|---|---|---|---|---|---|
| $Pr\{\text{Ext}\}$: | $\frac{1}{2}$ | $\frac{1}{2}$ | $\frac{1}{4}$ | $\frac{1}{10}$ | $\frac{1}{2}$ |
| $Pr\{\text{Cyt}\}$: | $\frac{3}{4}$ | $\frac{3}{4}$ | $\frac{1}{2}$ | $\frac{1}{4}$ | $\frac{3}{4}$ |
| $Pr\{\text{Nuc}\}$: | $\frac{3}{4}$ | $\frac{1}{4}$ | $\frac{1}{4}$ | $\frac{1}{8}$ | $\frac{1}{4}$ |
| | $v_1$ | $v_2$ | $v_3$ | $v_4$ | $v_5$ |
| *ExtMem* | $\frac{1}{2}$ | $\frac{1}{4}$ | $\frac{1}{16}$ | $\frac{1}{160}$ | $\frac{1}{320}$ |
| *Cytosol* | $-\infty$ | $\frac{3}{16}$ | $\frac{3}{32}$ | $\frac{3}{128}$ | $\frac{9}{512}$ |
| *Nucleus* | $-\infty$ | $-\infty$ | $\frac{3}{128}$ | $\frac{3}{1024}$ | $\mathbf{\frac{9}{2048}}$ |

#### 2 Incorporating the Mitochondria Compartment in the Signaling Model

The mitochondria compartment, denoted here as *Mt*, was added to the signaling model as an intermediate compartment, like the *Cytosol*, between the two terminal compartments *ExtMem* and *Nucleus*. S.Figure 1 shows a simplified diagram for the relationships among the different signaling compartments. We can reach a protein in the *ExtMem* only from another protein in the *ExtMem*. We can reach a protein in the *Cytosol* from another protein in one of the three compartments: *ExtMem*, *Cytosol*, or *Mt*. We can reach a protein in the *Mt* from another protein in one of the three compartments: *ExtMem*, *Cytosol*, or *Mt*. Finally, we can end up with a protein in the *Nucleus* from a protein in one of the three compartments: *Cytosol*, *Mt*, or *Nucleus*. This signaling model allows for having cyclic paths because of having two intermediate compartments: *Cytosol* and *Mt*. To adhere to the signaling assumptions outlined in the Methods subsection "Signaling Flow Structure and Assumptions" in the main manuscript, a path has to start with a protein in the *ExtMem*, end with a protein in the *Nucleus*, and have at least one protein in one of the intermediate compartments. So it is not necessary for a single path to have proteins in all the four cellular compartments, and consequently a path may have proteins in either three or four cellular compartments.

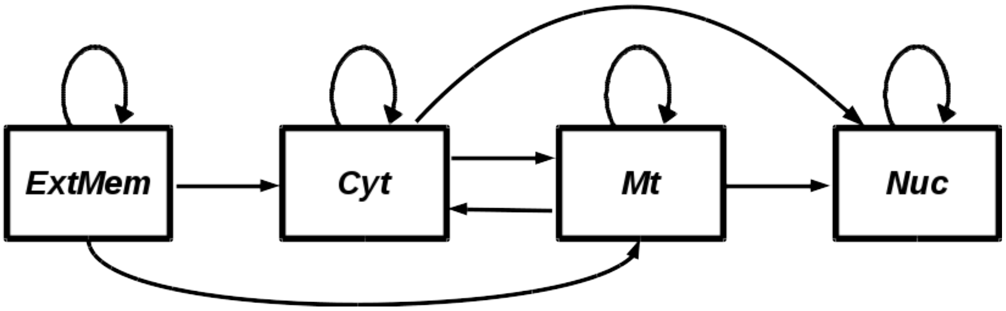


S.Figure 1. Signaling model when incorporating the mitochondria compartment.

Equations (1-7) are re-written here to consider the mitochondria compartment. The path final signaling score is computed as:

$$s(v_m, nuc) = \min\Big[s(v_{m-1}, cyt) + \mathcal{T}^{cyt}_{v_m}, s(v_{m-1}, cyt) + \mathcal{T}^{nuc}_{v_{m-1}}, s(v_{m-1}, Mt) + \mathcal{T}^{Mt}_{v_m}, s(v_{m-1}, Mt) + \mathcal{T}^{nuc}_{v_{m-1}}, s(v_{m-1}, nuc)\Big] + \mathcal{T}^{nuc}_{v_m}. \tag{15}$$

At node $v_j$, $j = 2, 3, \ldots, (m-1)$, the series of equations for the scores are:

$$s(v_j, ext) = s(v_{j-1}, ext) + \mathcal{T}_{v_j}^{ext} \tag{16}$$

$$\begin{aligned} s(v_j, cyt) = \min \Big[ & s(v_{j-1}, ext) + \mathcal{T}_{v_j}^{ext}, s(v_{j-1}, ext) + \mathcal{T}_{v_{j-1}}^{cyt}, \\ & s(v_{j-1}, Mt) + \mathcal{T}_{v_j}^{Mt}, s(v_{j-1}, Mt) + \mathcal{T}_{v_{j-1}}^{cyt}, s(v_{j-1}, cyt) \Big] + \mathcal{T}_{v_j}^{cyt}. \end{aligned} \tag{17}$$

$$\begin{aligned} s(v_j, Mt) = \min \Big[ & s(v_{j-1}, ext) + \mathcal{T}_{v_j}^{ext}, s(v_{j-1}, ext) + \mathcal{T}_{v_{j-1}}^{Mt}, \\ & s(v_{j-1}, cyt) + \mathcal{T}_{v_j}^{cyt}, s(v_{j-1}, cyt) + \mathcal{T}_{v_{j-1}}^{Mt}, s(v_{j-1}, Mt) \Big] + \mathcal{T}_{v_j}^{Mt}. \end{aligned} \tag{18}$$

$$\begin{aligned} s(v_j, nuc) = \min \Big[ & s(v_{j-1}, cyt) + \mathcal{T}_{v_j}^{cyt}, s(v_{j-1}, cyt) + \mathcal{T}_{v_{j-1}}^{nuc}, \\ & s(v_{j-1}, Mt) + \mathcal{T}_{v_j}^{Mt}, s(v_{j-1}, Mt) + \mathcal{T}_{v_{j-1}}^{nuc}, s(v_{j-1}, nuc) \Big] + \mathcal{T}_{v_j}^{nuc}. \end{aligned} \tag{19}$$

The base case for these recurrence relations are:

$$s(v_1, ext) = \mathcal{T}_{v_1}^{ext} \tag{20}$$
$$s(v_1, cyt) = \infty \tag{21}$$
$$s(v_1, Mt) = \infty \tag{22}$$
$$s(v_1, nuc) = \infty. \tag{23}$$

### 3 Evaluation of Multiple Pathways

We start by defining the precision and recall for individual pathways, and then extend this to the case of multiple pathways—the aggregate pathways. For each pathway, we compute its precision and recall (PR) values using its set of positives $P$, its set of negatives $N$, and its set of predicted interactions $X$. The interactions in $X$ are ranked by the number/rank of the first path an interaction appears in (ascending order). Let $X_i$ denote the set of unique predicted interactions up to the path $i$. The precision and recall for $X_i$ are computed as:

$$Precision_i = \frac{|X_i \cap P|}{|X_i|} \quad and \quad Recall_i = \frac{|X_i \cap P|}{|P|}, \tag{24}$$

where $|S|$ means the number of elements in set $S$.

For the case of computing the PR values for $m$ pathways $p_1, p_2, \ldots, p_m$, we have $m$ distinct collections of positive interactions, negative interactions, and ranked predicted interactions, denoted as $P^j$, $N^j$, and $X^j$ for the $j$th pathway, respectively. We aggregate the sets of the ranked predictions as:

$$X = \bigcup_{j=1}^{m} \left[(e,k) \ \text{ for } \ e,k \in X^j\right], \tag{25}$$

where $e$ is a predicted interaction and $k$ is its rank in $X^j$. We then rank the elements in $X$ by the value $k$. Similarly, we aggregate the sets of positives and negatives as:

$$P = \bigcup_{j=1}^{m} \left[p \ \text{ for } \ p \in P^j\right] \quad and \quad N = \bigcup_{j=1}^{m} \left[n \ \text{ for } \ n \in N^j\right]. \tag{26}$$

We used these three aggregate sets, $X$, $P$, and $N$, to compute the precision and recall values using Equation (24). The size of the negatives set is 25 times the size of the positives set for the individual pathways.

#### 4 The Color-Coding (CC) Technique

For a weighted, directed graph $G = (V, E)$, where $V$ is the vertices set and $E$ is the directed edges set, each edge $(u, v) \in E$ has a weight $w_{uv} \in [0, 1]$. The color_coding (CC) algorithm [1] can be used to compute simple paths, such that each starts at a specific vertex and ends at another specific vertex, and no vetex is visited more than once. Given a graph $G$, a set $R$ of a path starting points (e.g. cellular membrane receptors) and a set $T$ of ending points (e.g. transcriptional regulators (TRs)), and a fixed number $q$ representing the path length (number of vertices), CC computes an optimal/sub-optimal solution for the problem of finding a path with the minimum reconstruction cost $r_i$, where the path $P_i = (v_1, v_2, \ldots, v_q)$ is comprised of $q$ vertices that begin at a receptor ($v_1 \in R$) and end at a TR ($v_q \in T$). Note that the shortest path is the one whose edge weights product is the highest among all paths since we take the negative log-transform of the edge weights at the reconstruction step.

The CC method can be integrated with Yen's algorithm to compute a ranked list of the $k$ shortest paths $\mathcal{P} = \langle P_1, P_2, \ldots, P_k \rangle$. Each path $P_i$ is ranked by its reconstruction cost $r_i$, and $r_i \leq r_{i+1}$ for every $i$. We will describe the CC algorithm first and then will illustrate how to integrate CC with Yen's algorithm.

##### 4.1 The Original CC Algorithm

For a fixed length $q$, CC randomly assigns to each vertex in the graph a uniformly distributed color (label) from $\{1, 2, \ldots, q\}$, and then finds a *colorful* path such that each one of the $q$ vertices composing the path has a distinct color. To guarantee that each path starts at a vertex in $R$ and terminates at a vertex in $T$, we add to $V$ a new vertex $s$, a *super source*, and link it to every vertex in $R$ with edge weight $w_{sv} = 0, \forall v \in R$, and add to $V$ another vertex $t$, a *super target*, and link it to every vertex in $T$ with edge weight $w_{vt} = 0, \forall v \in T$. Hence, a colorful path has to start at $s$ and ends at $t$ and will be of length $q' = q + 2$. Instead of randomly assigning a color to each vertex, we assign the color 0 to $s$ and the color $(q + 1)$ to $t$.

CC uses a dynamic programming approach to compute the minimum-weight colorful path. Let the function $c(v)$ returns the color $c$ of a node $v$ in the graph. Let $C_j$ be a set of $j$ distinct colors such that $C_j \subseteq \{0, 1, \ldots, q, q+1\}$. Define the function $W(v, C_j)$ to be the minimum weight of a simple path that starts at $s$ (with color 0), ends at $v$, is of length $|C_j| = j + 1$ ($j$ vertices in addition to $s$), and visits one vertex of each color in $C_j$. If no such path exists, then $W(v, C) = \infty$.

The initial values are $W(s, C_0) = 0$, where $C_0 = \{0\}$. For every value of the monotonically increasing index $j : j = 1, 2, \ldots, q, q + 1$, the dynamic program uses the following recurrence to compute $W(v, C_j)$ for every $v \in V$:

$$W(v, C_j) = \min_{\substack{\{u,v\} \in E, \text{ where} \\ c(u) \in C_{j-1}, \text{ and} \\ C_{j-1} = C_j \setminus c(v)}} \Big( W(u, C_{j-1}) + w_{u,v} \Big) \tag{27}$$

That is, the dynamic checks all incoming edges to $v$ where the tail is in $C_{j-1}$ and $C_j$ differs from $C_{j-1}$ by exactly $c(v)$. The dynamic program terminates after computing $W(t, C_{q+1})$, where $C_{q+1} = \{0, 1, \ldots, q, q+1\}$ with any order of the colors, except for the start, 0, and the end, $q + 1$. This represents finding a simple path of length $(q + 2)$. By excluding the two vertices $s$ and $t$, the path will have a length of $q$, which is the solution to the problem.

The constraint of a colorful path (distinct colors of the path vertices) ensures that the reconstructed path is simple. The random designation of colors to the vertices leads to an optimal/sub-optimal solution, if one exists. So, a large number of iterations is required to increase the probability of finding a colorful path. For a path of length $q$, we have $q^q$ ways to color the path, and have $q!$ ways to make a colorful path, i.e. no repeated colors per path. So, the probability of having a colorful path is:

$$P\{\text{colorful path}\} = \frac{q!}{q^q}.$$

The probability of failure, i.e. not having a colorful path, is:

$$P\{\text{failure}\} = 1 - \frac{q!}{q^q}.$$

The probability of failure is essentially larger than the probability of success. Hence, the CC algorithm is repeated a very large number of iterations to increase the probability of having a colorful path. For a $d$ number of iterations, the probability of failure will be:

$$P\{\text{failure}\} = \left(1 - \frac{q!}{q^q}\right)^d.$$

If we up limit the probability of failure to $\epsilon$, such as 0.001 in this study, we can compute $d$ as:

$$d = \frac{\log \epsilon}{\log(1 - \frac{q!}{q^q})}.$$

The number of iterations, $d$, increases exponentially with increasing the path length, $q$, and/or increasing the probability of success (i.e. decreasing $\epsilon$) [1].

#### 4.2 Extending the CC Technique by Integrating the proteins Cellular Information

To incorporate cellular localization information about proteins, we follow the method in [2]. Since we only consider intracellular signaling that begins with activation of a membrane-bound protein receptor and is transmitted to a nucleus DNA-binding transcription factor through PPIs within the cytosol, we focus on three cellular compartments: a combination of extracellular fluid and cell membrane (*ExtMem*), which represents where a receptor may be located, *Cytosol*, and *Nucleus*. We define the localization function $L(v) : L(v) \subseteq \{ExtMem, Cyt, Nuc\}$ that returns the possible compartments a protein could be found within. We rely on the ComPPI dataset [3] to assign cellular compartments to each protein: $L(v) = c \iff Pr($ protein $v$ can be found in compartment $c) > 0$. We can only reach a protein in *ExtMem* from another protein in *ExtMem*, we can reach a protein in *Cytosol* from another protein in either *ExtMem* or *Cytosol*, and we can reach a protein in *Nucleus* from another one in either *Cytosol* or *Nucleus*.

The above recurrence in Equation (27) needs to be modified to integrate the localization information. The modified version is:

$$W(v, C_j) = \min_{\substack{\{u,v\} \in E \\ c(u) \in C_{j-1} \\ C_{j-1} = C_j \setminus c(v)}} \left\{ \Big( W(u, C_{j-1}) + w_{u,v} \Big) \cdot F(u,v) \right\},$$

where $F(u, v)$ is a function in terms of the localization of the two proteins of the edge $(u, v)$ and is defined as:

$$F(u,v) = \begin{cases} 1 & \begin{cases} u = s, v \in R, ExtMem \in L(v), \\ ExtMem \in L(u), ExtMem \in L(v) \text{ or } Cyt \in L(v), v \notin T, \\ Cyt \in L(u), Cyt \in L(v) \text{ or } Nuc \in L(v), u \notin R, \\ Cyt \in L(u) \text{ or } Nuc \in L(u), Nuc \in L(v), u \notin R, \\ u \in T, Nuc \in L(u), v = t, \end{cases} \\ 0 & ; otherwise. \end{cases}$$

$W(t, C_{l+1})$ in this case represents a simple path of length $(l + 2)$ that, after excluding the two vertices $s$ and $t$, preserves the signaling hierarchy by starting at the extracellular fluid or at the cell membrane and ending inside the nucleus.

#### 4.3 Yen's Algorithm

If we need to generate a $k$-paths list, e.g. $k = 20,000$ as in this study, we need to run CC a number of iterations greatly larger than $k$ to account for the trials of non-colorful paths. This can take up to days, if not weeks, for a single pathway if the interactions network is very large. So, we augment CC with Yen's algorithm [4] to compute the $k$-shortest paths based on the CC method. We call this the *Yen_CC* method. A formal description of Yen's algorithm is immediately following.

Given a weighted, directed graph $G = (V, E)$, where $V$ is the set of the vertices, $E$ is the set of the directed edges, and each edge $(u, v) \in E$ has a weight $w_{uv} \in [0, 1]$, and given two vertices $s$ and $t$ in $V$, Yen's algorithm uses any shortest path algorithm, such as Dijkstra's algorithm, as a subroutine to find the $k$ shortest loopless paths from $s$ to $t$. The shortest path subroutine is first employed to compute a single path that is the shortest one from $s$ to $t$ in $G$, and after that Yen's algorithm takes place to find the second shortest path and so on. In general, let the $i$th shortest $s$-$t$ path in $G$ be $\pi_i$ and let the $j$th vertex in that path be $\pi_{i,j}$. Yen's algorithm operates on the principle that each new shortest path $\pi_i$ can be generated from some previous shortest path $\pi_{i'}$, $i^{'} < i$, by assuming that $\pi_i$ deviates from $\pi_{i'}$ after some vertex $\pi_{i',j'}$. Yen's algorithm computes this path by executing a shortest path search from $\pi_{i',j'}$ to $t$ on a graph $G^{'}$, which is constructed by removing from $G$ all the vertices in $\{\pi_{i',1}, \pi_{i',2}, \ldots, \pi_{i',j'-1}\}$ in addition to any outgoing edges from $\pi_{i',j'}$, which are in a previously found path. This construction guarantees that the path found in $G^{'}$ represents a new, loopless $s$-$t$ path (Supplementary material of [5]). In simple words, once Yen's algorithm finds a path, it searches for alternative paths that differ from the discovered path in one or more edges, i.e. it searches for new partial paths. In *Yen_CC*, we employ the CC method as the shortest path subroutine. Hence, in *Yen_CC*, instead of running a new iteration to find a complete colorful path, the iteration will look for a partial colorful path, leading to reduction in the search space and time. Since *Yen_CC* takes as an input a single path length value, we run *Yen_CC* across a sequence of path lengths, combine all the paths in a single list, and then re-order them all based on the reconstruction cost. For weighted graphs, paths with more vertices do not imply having higher reconstruction weights, that is why we re-order the concatenated paths in the final list.

### 5 Signaling Pathways

We used a set of four NetPath pathways [6] to evaluate the proposed method. S.Table 1 summaries the number of protein-protein interactions (PPIs), receptors, transcription regulators (TRs) for each pathway.

S.Table 1. Signaling pathways used in this study and numbers of their interactions, receptors, and transcription regulators (TRs) for both the $PLNet_2$ and HIPPIE [7] interactomes and for the condition of the complete interactome and the condition of the interactome intersected with the ComPPI database [3].

| Pathways | Interactome Condition | $PLNet_2$ | | | HIPPIE Interactome | | |
|---|---|---|---|---|---|---|---|
| | | PPIs | Receptors | TRs | PPIs | Receptors | TRs |
| $\alpha 6 \beta 4$ Integrin | Complete | 192 | 7 | 3 | 184 | 7 | 3 |
| | $\cap$ ComPPI | 115 | 7 | 3 | 107 | 6 | 3 |
| EGFR1 | Complete | 1308 | 6 | 33 | 1274 | 6 | 32 |
| | $\cap$ ComPPI | 659 | 6 | 33 | 627 | 4 | 31 |
| IL2 | Complete | 199 | 3 | 12 | 178 | 3 | 12 |
| | $\cap$ ComPPI | 171 | 3 | 12 | 150 | 3 | 12 |
| Wnt | Complete | 347 | 14 | 14 | 346 | 14 | 14 |
| | $\cap$ ComPPI | 168 | 14 | 14 | 166 | 11 | 12 |

### 6 Protein Compartments

S.Table 2. Protein compartment information in $PLNet_2$.

| Cellular Compartment | # Proteins with a single compartment | # Proteins with multiple compartments | Average localization score |
|---|---|---|---|
| *ExtMem* | 17 | 9,993 | 0.8214 |
| *Cytosol* | 464 | 12,726 | 0.8595 |
| *Nucleus* | 923 | 10,461 | 0.8824 |
| *Mitochondria* | 78 | 2,718 | 0.7942 |
| *Secretory* | 25 | 6,034 | 0.7759 |

### 7 Number of Ties

S.Table 3 reports the number of path groups that share the same reconstruction score after applying PathLinker (*PL*) on the original $PLNet_2$ interactome and the filtered interactome using the cellular localization information. Filtering the interactome by keeping only the spatially coherent interactions reduced the number of ties in all the pathways. However, ties still dominate the reconstructions, and this urges the need for a way for breaking these ties.

S.Table 3. The number of ties (path groups sharing the same reconstruction score) for *PL* applied on the original $PLNet_2$ interactome and the filtered interactome.

| Pathway | Original Interactome | Filtered Interactome |
|---|---|---|
| $\alpha 6 \beta 4$ Integrin | 82 | 58 |
| EGFR1 | 17 | 15 |
| IL2 | 28 | 17 |
| Wnt | 43 | 22 |

### 8 Pathway Reconstructions for $\alpha6\beta4$ Integrin, EGFR1, IL2, and Wnt

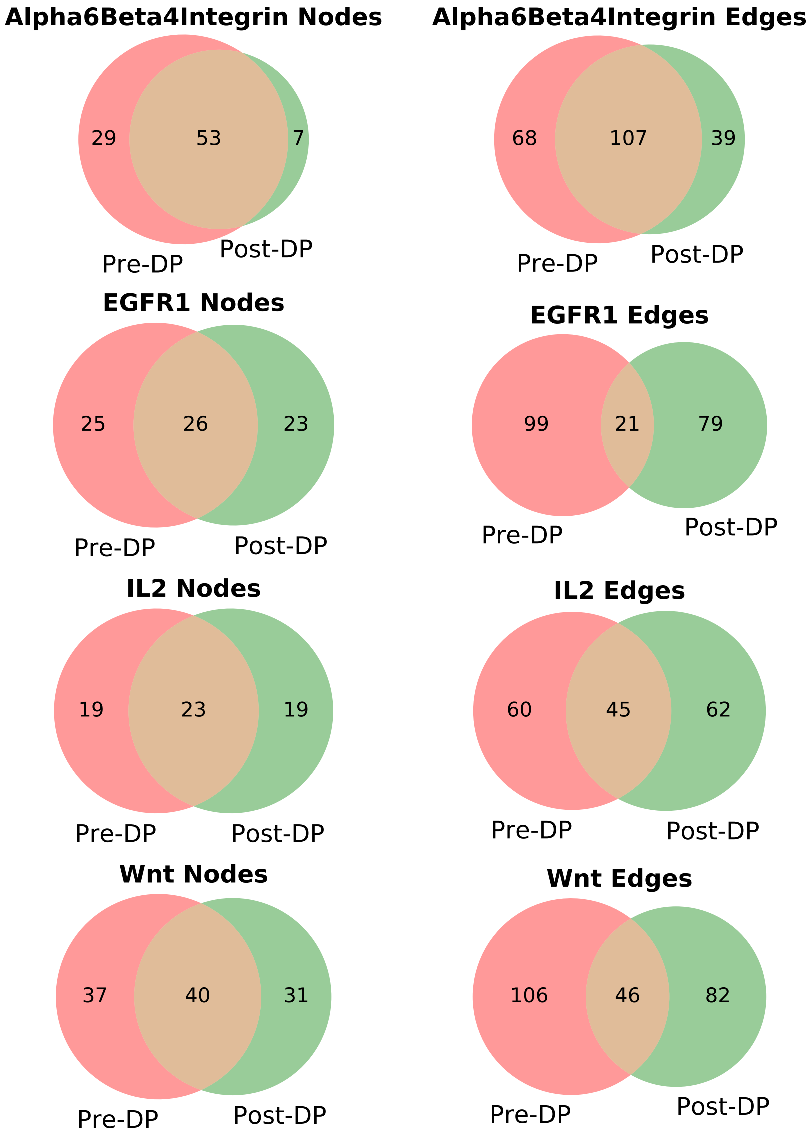


S.Figure 2. Number of nodes and edges for the first 100 paths in each pathway reconstruction before (Pre-DP) and after (Post-DP) applying the dynamic program.

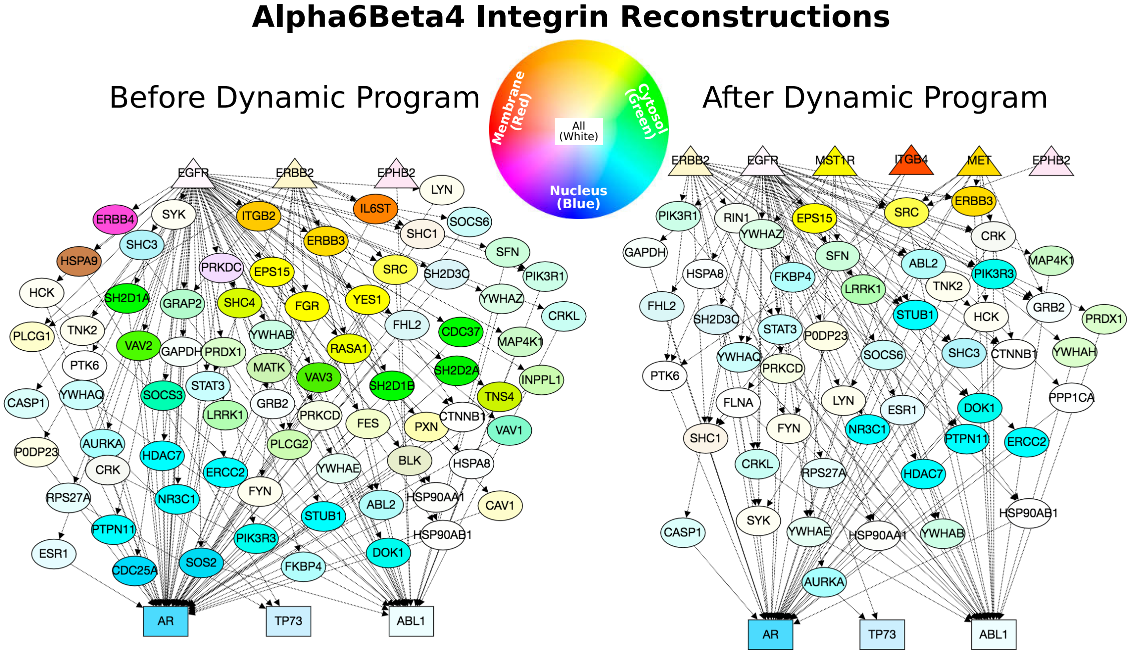


S.Figure 3. *LocPL* pathway reconstructions (first 100 paths) for $\alpha6\beta4$ Integrin before applying the dynamic program (left) compared to after applying the dynamic program (right). Receptors are labeled as triangles, transcriptional regulators are rectangles, intermediary proteins are ellipses. Color denotes compartment localization; proteins may belong to multiple compartments (and will be lighter shades). Networks generated using GraphSpace [8] and are available at http://graphspace.org/graphs/?query=tags:LocPL.

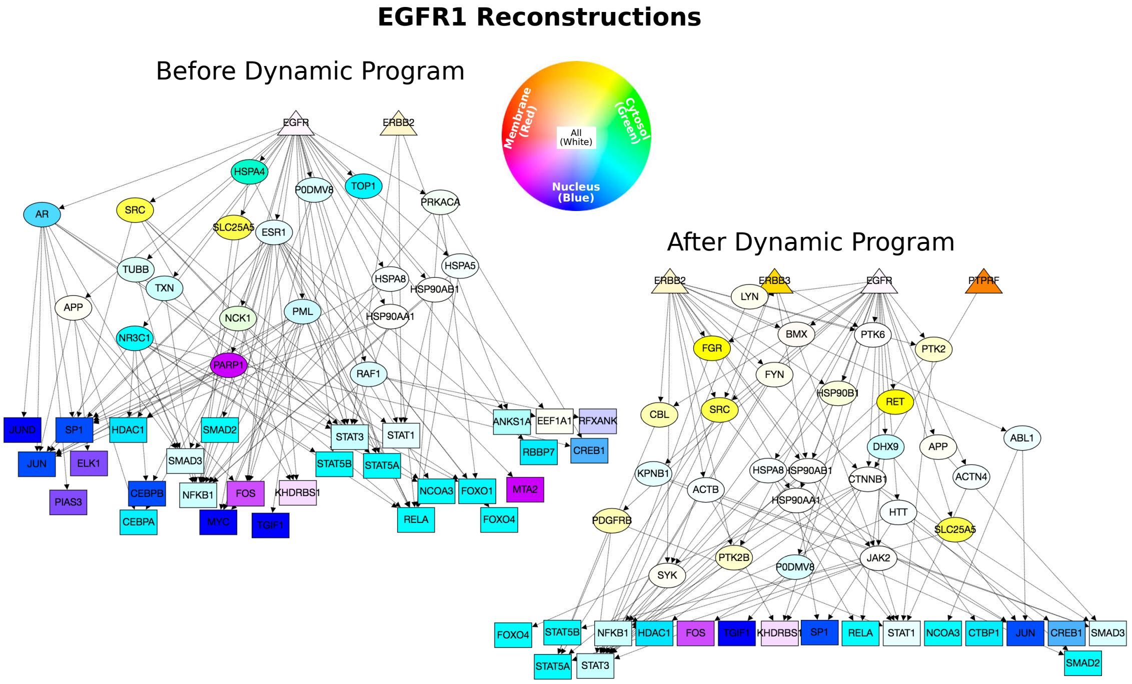


S.Figure 4. *LocPL* pathway reconstructions (first 100 paths) for EGFR1 before (left) and after (right) the dynamic program. Nodes are described as in S.Figure 3 and are available at http://graphspace.org/graphs/?query=tags:LocPL.

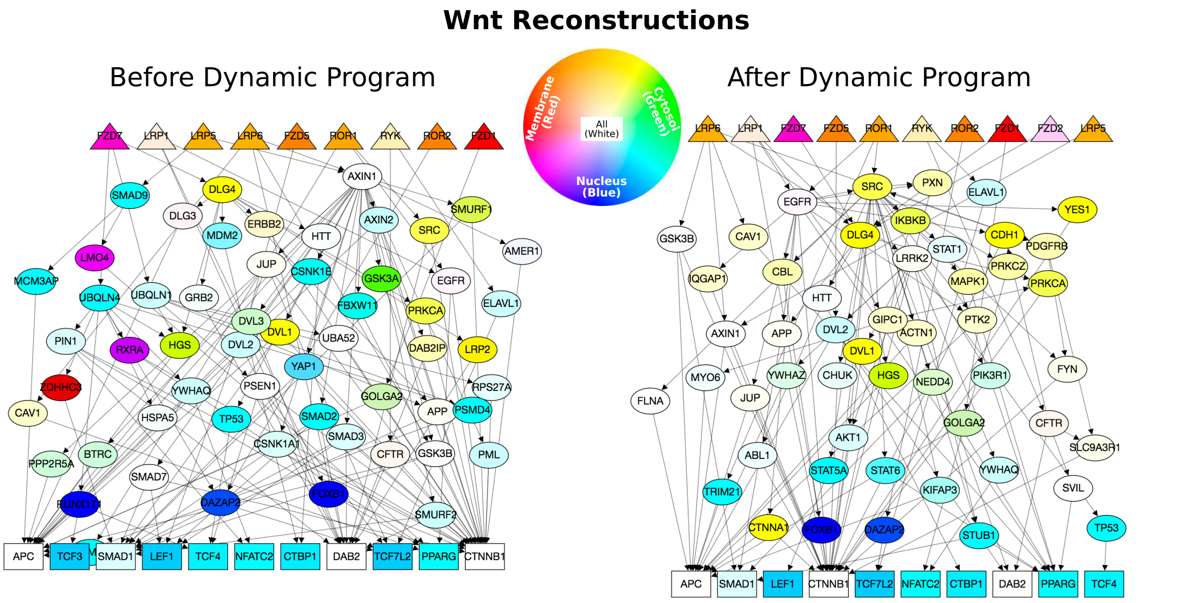


S.Figure 5. *LocPL* pathway reconstructions (first 100 paths) for Wnt before (left) and after (right) the dynamic program. Nodes are described as in S.Figure 3 and are available at http://graphspace.org/graphs/?query=tags:LocPL.
